# Efficient and multiplexed tracking of single cells using whole-body PET/CT

**DOI:** 10.1101/2023.08.23.554536

**Authors:** Hieu T.M. Nguyen, Neeladrisingha Das, Yuting Wang, Carlos Ruvalcaba, Brahim Mehadji, Emilie Roncali, Charles K.F. Chan, Guillem Pratx

## Abstract

*In vivo* molecular imaging tools are crucially important for elucidating how cells move through complex biological systems, however, achieving single-cell sensitivity over the entire body remains challenging. Here, we report a highly sensitive and multiplexed approach for tracking upwards of 20 single cells simultaneously in the same subject using positron emission tomography (PET). The method relies on a new tracking algorithm (PEPT-EM) to push the cellular detection threshold to below 4 Bq/cell, and a streamlined workflow to reliably label single cells with over 50 Bq/cell of ^18^F-fluorodeoxyglucose (FDG). To demonstrate the potential of method, we tracked the fate of over 70 melanoma cells after intracardiac injection and found they primarily arrested in the small capillaries of the pulmonary, musculoskeletal, and digestive organ systems. This study bolsters the evolving potential of PET in offering unmatched insights into the earliest phases of cell trafficking in physiological and pathological processes and in cell-based therapies.

**TEASER:** A novel PET imaging workflow reveals the fate of single cells across the body.

## 1. Introduction

Over the recent decades, significant progress has been made in the field of molecular imaging to develop techniques specifically aimed at tracking the movement of cells inside the body. These techniques have found application in a range of different biological fields, where they have been used to characterize the spread of tumor cells and subsequent formation of metastases,^1^ the spatial kinetics of immune cells in response to pathogens,^2,3^ the migration of regenerative stem cells towards damaged tissues,^4–6^ and other key biological processes such as embryonic development and immune cell patrolling.^7,8^ These imaging tools have become crucial for monitoring the fate of cells as they move across the body and understanding the role that this migration plays in the context of physiological and pathological processes and cell-based therapies.

Positron emission tomography (PET), known for its exceptional sensitivity across the entire body, presents a compelling platform for tracking cellular dynamics in both human subjects and preclinical animal models. ^9–11^ However, applications of this technique have been predominantly conducted on a population level, which is incompatible with the need to elucidate the trajectories of individual cells. To address this limitation, we previously devised a methodology to expand the capabilities of PET into the realm of single-cell imaging. Using mesoporous silica nanoparticles (MSNs), we efficiently labeled cells with ^68^Ga and visualized the migration and arrest of solitary breast cancer cells in lung capillaries.^12^ Cells containing upwards of 27 Bq/cell produced sufficient signal to be clearly distinguished, and sensitivity was further improved by the use of a novel tracking algorithm called CellGPS.^12–14^ These results firmly established PET as a novel method for tracking single cells across the body.

However, the promise that PET holds for tracking single cells can only be realized once remaining challenges have been addressed. Foremost among these is the lack of cellular multiplexing capabilities in the initial implementation of this technique, which could track just one cell per animal. Because of this low throughput, it was prohibitive to conduct larger studies involving 10 or more cells. Secondly, despite its high efficiency, the cell labeling workflow based on MSNs proved laborious and at times inconsistent due to its complexity and lack of automation. This limitation became particularly restrictive as we sought to deploy this technique on a more routine basis in the context of studying the fate of cancer cells during metastasis. Thirdly, a more fundamental question arises concerning the lower limit of detection achievable with PET. While we could previously image single cells containing around 20 Bq, theoretical considerations suggest that the detection threshold could be lowered even further, which would have implications for tracking different types of cells and for improving the temporal resolution.

We here report a multiplexed approach for tracking upwards of 20 single cells concurrently in the same subject using PET. The proposed workflow represents a considerable improvement in nearly every aspect, including cell labeling, single-cell dispensing, data acquisition, data reconstruction, and disease modeling. In addition to the 20-fold improvement in multiplexing capability, we demonstrate a detection threshold of 4 Bq for single cells, a five-fold improvement in sensitivity. The entire methodology, while solely relying on commercially available reagents and instruments, could nonetheless achieve high labeling efficiency in the range of 50-100 Bq/cell. Finally, the measured single-cell PET signal only arises from live cells, which bolsters the specificity of the assay. Collectively, these advances will facilitate widespread adoption of this technique across institutions while providing a reliable, straightforward, and accurate method for tracking multiple cells in vivo in crucial biological experiments.

## 2. Results

### A streamlined workflow for cell radiolabeling and dispensing

A crucial step towards the routine use of PET for tracking single cells *in vivo* is the design of an efficient and straightforward workflow for radiolabeling and dispensing single cells (**Figure 1**). The key features of this new method are that (*i*) labeling is achieved through a commonly available PET tracer, ^18^F-fluorodeoxyglucose (FDG), and (*ii*) single cells are dispensed using an automated microfluidics device. Compared to other radiolabeling techniques, FDG can be inexpensively sourced from commercial vendors and central facilities without requiring specialized radiochemistry; hence, the method can be easily adopted by other researchers. Another advantage of FDG over ^68^Ga-MSN lies in its longer half-life (2 hours) and shorter positron range, which translates into more accurate localization of single cells over longer periods of time. In addition, the use of an automated single-cell dispenser shortens the time required for isolating and dispensing viable single cells by eliminating the laborious steps involved in limiting cell dilution and microscopic verification. We further enhanced the detectability of single cells using a dual-layer BGO/LYSO PET scanner^15,16^ with high sensitivity (∼10.4%) and resolution (∼1.2 mm). Notably, the use of BGO as a scintillator material in this scanner results in a lower rate of background coincidence events compared to lutetium-based scintillators, which is a crucial advantage when imaging weakly radioactive cells. These background events are caused by the natural occurrence of radioactive ^176^Lu in the scintillator material.^17,18^

**Figure 1.**
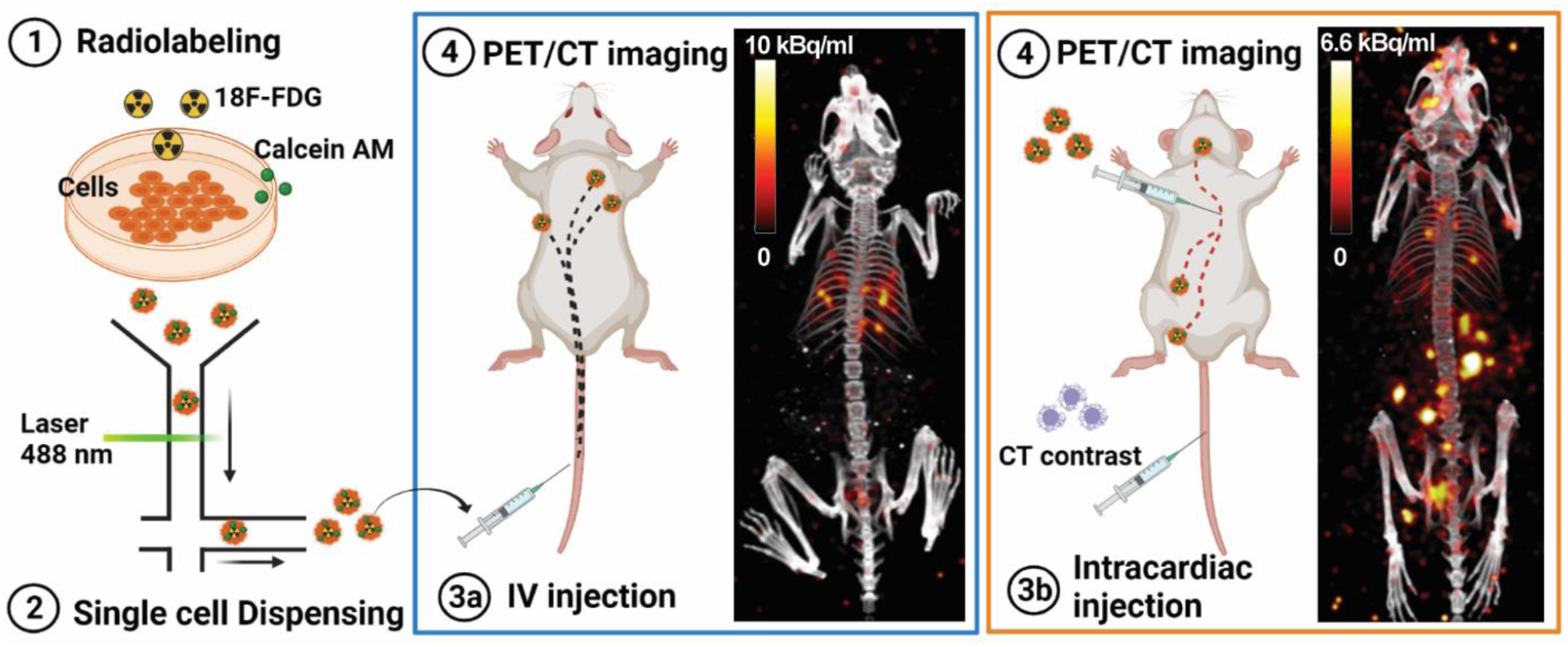
Workflow for tracking single radiolabeled cells *in vivo* with PET. Cancer cells (B16) were radiolabeled with FDG (1), then sorted and dispensed as single cells using a microfluidic device (2). As a proof of concept, the labeled cells were injected into a murine model via either intravenous (3a) or intracardiac (3b) routes. Cells injected intravenously were trapped in the lungs, whereas those injected intracardially were widely disseminated throughout the entire body. We used PET/CT and a custom algorithm to estimate the 3D locations of these single cells (4).

To achieve high FDG labeling, cancer cells were incubated with FDG (370 MBq/ml) for 1 h after a “fasting” period of 1 h in glucose-free media. After removing residual FDG, single FDG-labeled cells were dispensed into microcentrifuge tubes using a microfluidic-based cell sorter. Consistent with our previous study,^12^ these single cells appeared as distinct focal spots in PET images (Fig. 2a**, Supplementary** Fig. 1). To demonstrate that these focal spots were induced by single cells, we partitioned the contents of each vial into two or three new vials and imaged the samples a second time (**Supplementary** Fig. 1a). We also used gamma counting to measure the radioactivity of the vial before and after partitioning the samples (**Supplementary** Fig. 2). In nearly all experiments, we observed that the radioactivity remained confined to a single vial after partitioning, indicating that the vials initially contained single cells, which could not be split among multiple vials. In rare cases (<5%), however, we observed radioactivity in two or more vials after partitioning, which we attributed to the presence of multiple cells in the sample or the release of the radioactivity into the medium following the death of a labeled cell. To further investigate the latter scenario, we used PET to image cells that were deliberately killed by adding lysis buffer. In these samples, the PET signal was not confined to a localized focal point but was instead distributed within the entire vial (Fig. 2b**, Supplementary** Fig. 1b). Thus, since FDG diffuses out of the cell upon its death, the presence of a focal signal in the PET image specifically reflects the viability of the imaged cells. Collectively, these *in vitro* results demonstrate the benefits of combining FDG labeling with microfluidic cell dispensing to efficiently and reliably visualize multiple cells by PET imaging.

**Figure 2.**
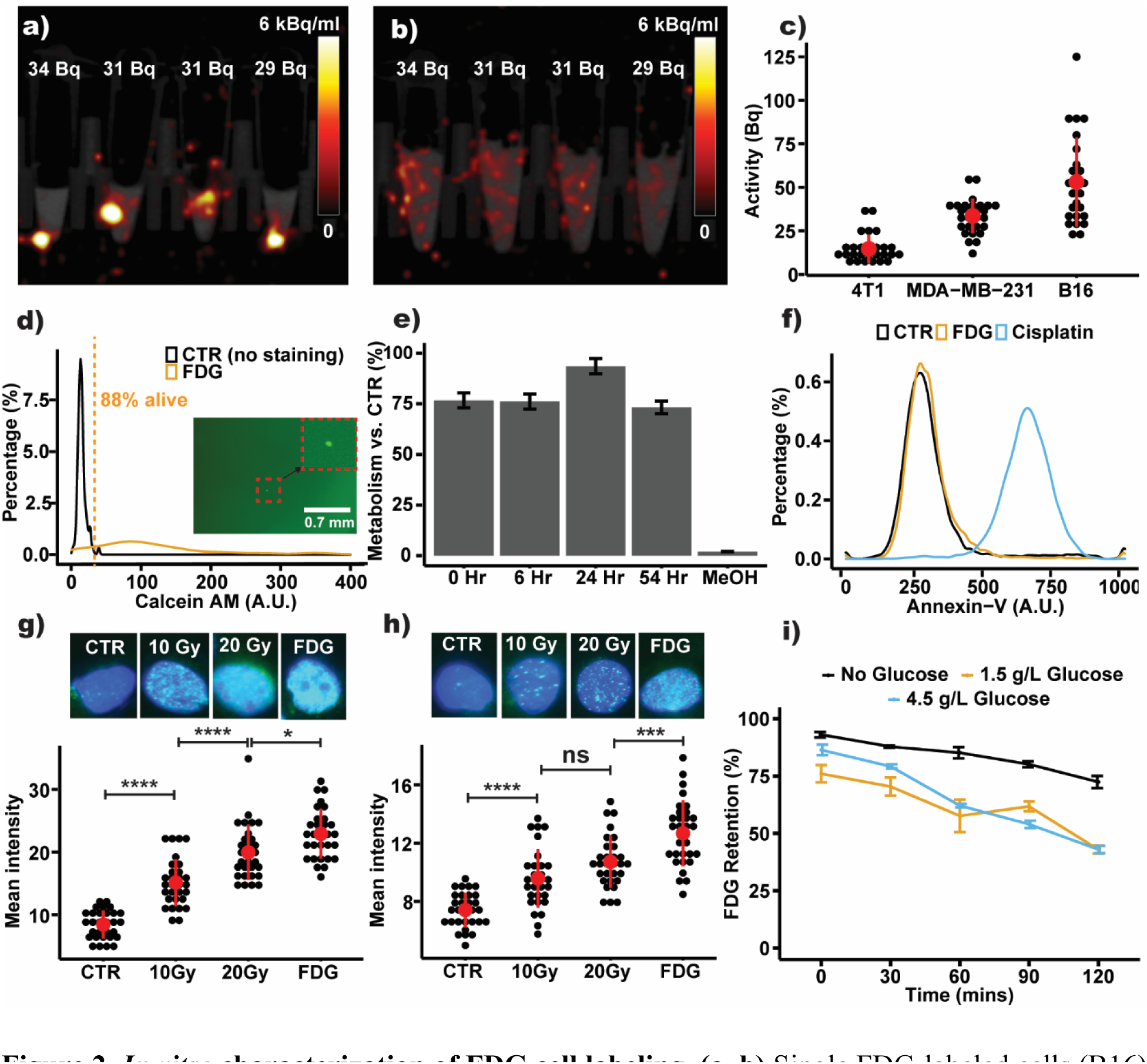
*In vitro* characterization of FDG cell labeling. **(a, b)** Single FDG-labeled cells (B16) were placed in vials and imaged with PET/CT before and after adding lysis buffer, showing that focal PET signal is a characteristic feature of live cells. **(c)** The radioactivity of single cells was measured with a gamma counter, revealing that B16 cells could be labeled with FDG more effectively than MDA-MB-231 and 4T1 cells. **(d)** Viability of FDG-labeled cells was >88%, as assessed by Calcein-AM staining. Inset picture: fluorescence micrograph confirming that the device dispensed exactly one cell. **(e)** CCK-8 assay, demonstrating a 75% cell metabolic rate 54 hours after labeling, relative to control cells. **(f)** Annexin-V assay, showing comparable levels of cell apoptosis in labeled and control cells 3 hours after labeling and significantly less than the positive control (Cisplatin)**. (g, h)** DNA damage characterized using γH2AX staining for control, X-ray irradiated, and FDG-labeled cells, measured 10 minutes (g) and 24 hours after labeling (h). **(i)** FDG efflux from FDG-labeled cells in the presence of different concentrations of D-glucose (0, 1.5g/L, and 4.5g/L).

The labeling efficiency was further characterized by single-cell gamma counting for three cell lines (Fig. 2c). The average FDG uptake was highest for B16 cells (52 Bq/cell), followed by MDA-MB-231 (32 Bq/cell) and 4T1 cells (13 Bq/cell). The observed difference in FDG uptake was likely due to intrinsic differences in basal glucose metabolism. Moreover, this metric was reproducible and did not vary significantly across experiments (data not shown). Compared to the labeling method based on ^68^Ga-MSNs reported by Jung et al., where average activity for MDA-MB-231 labeled cells varied between 3-10 Bq/cell batch-to-batch,^12^ the labeling protocol using FDG was more consistent and yielded higher average activity within the same cell line. Additionally, the radioactivity of single cells can be measured using PET by computing the average uptake within regions of interest (ROIs) centered on individual cells. The cell activity characterized through this method was consistent with the radioactivity measured using a gamma counter (**Supplementary** Fig. 3), thus suggesting that PET can quantitatively measure single-cell activity *in vivo*.

We also found that FDG uptake was strongly influenced by the concentration of glucose in the media. Though we labeled cells in glucose-free media, a small amount of glucose was present in the fetal bovine serum (FBS) supplement, which ended up significantly influencing the efficiency of the labeling procedure (**Supplementary** Fig. 4). Relative to standard 10% FBS media (12.5 mg/dL glucose), FDG uptake increased significantly when using 1% FBS media (1.25 mg/dL glucose). There was no significant difference when we further reduced glucose concentration using 10% dialyzed FBS (0.1 mg/dL glucose). The presence of FBS was necessary for the viability and normal metabolism of cells, as we observed reduced uptake when no FBS was added to the media. Based on these results, we selected 1% FBS as the standard condition for cell labeling.

In addition, we examined the viability and behavior of B16 cells labeled with FDG according to several assays. Exposure to ionizing radiation during the labeling procedure may interfere with the normal functions of the labeled cells. Calcein AM was used as a marker of cell viability during single-cell dispensing. Analysis of FDG-labeled cells revealed that 88% of them were positive for Calcein AM staining (gated above the autofluorescence signal of the control cells; Fig. 2d). To further assess the potential toxicity of the labeling procedure, metabolic activity was measured at times 0, 6, 24 and 54 hours using the colony-counting kit colorimetric assay (CCK-8). Compared to control cells, FDG-labeled cells maintained >75% metabolism for up to 54 hours post-labeling (Fig. 2e).

We further investigated the biological fate of labeled cells and whether they underwent apoptotic or necrotic cell death following exposure to radioactivity, using Annexin-V and Sytox Green as markers for these events. Three hours after FDG labeling, no significant increase in Annexin-V staining was observed relative to control cells. In contrast, cells treated with Cisplatin (positive control; 100 μM, 48 hours) demonstrated strong binding of the marker (Fig. 2f). The FDG-labeled cells also presented minimal Sytox signal, indicating that the cell membrane was intact following radiolabeling (**Supplementary** Fig. 5).

DNA damage commonly occurs following radiation exposure. To quantify this effect, we measured DNA double-strand break by γH2AX immunostaining. To provide a quantitative reference, the intensity of the γH2AX signal was also measured for cells exposed to 10 and 20 Gy of X-ray radiation (Fig. 2g). Taken together, these results suggest considerable DNA damage 10 min post-radiation, though the damage was substantially repaired 24 hours after radiolabeling (Fig. 2h). In our estimate, most of the DNA damage is incurred during the hour-long incubation with FDG. A simple calculation based on positron energy deposition suggests that the absorbed dose experienced by the cells during the 1 h incubation was approximately 44 Gy (Supplementary Method). Because positrons deposit their energy over a distance (∼1 mm) much longer than the typical cell diameter (0.01 mm), the radiation dose experienced by the labeled cells *in vivo* is only a small fraction (<1 %) of the total radiation dose.

To further characterize the behavior of FDG-labeled cells, we assessed the migration of B16 cells *in vitro* in response to a small mechanical scratch to the surface of the cell monolayer. Based on observations at 0, 6, 24, and 48 hours, we found that FDG-labeled cells could close the gap at a similar rate to control cells, whereas the gap expanded for cells treated with Cisplatin (positive control; 100 μM for 48 hours) during the observation period (**Supplementary** Fig. 6), thus suggesting that the labeling procedure did not affect the ability of the cells to migrate *in vitro*.

Finally, we studied the rate of FDG efflux from labeled B16 cells for media containing different glucose concentrations (0, 1.5, and 4.5 g/L D-glucose). Cellular efflux of the radiotracer can lower tracking accuracy by causing loss of cell signal and higher background radioactivity. At the 120 min time point, cells retained upwards of 75% of the initial FDG radioactivity in glucose-free media, compared to 48% when higher glucose concentrations (1.5 g/L and 4.5 g/L) were used (Fig. 2i). The influence of extracellular glucose on FDG efflux from the labeled cells is relevant considering the typical concentration of glucose in the blood of healthy fasting mice (0.8-1 g/L).^19^

In summary, high labeling radioactivity could be achieved with FDG, and most labeled cells remained viable throughout the time frame of a typical ^18^F tracking experiment (a few hours). While an increase in DNA damage was noted, the labeling procedure did not alter the ability of tumor cells to migrate *in vitro*. Efflux was increased in the presence of extracellular glucose, yet cells retained approximately half of the initial labeling radioactivity, which is comparable to other labeling strategies.

### *In vivo* imaging of multiple FDG-labeled individual cells with PET

Having characterized the *in vitro* efficacy and potential toxicity of FDG as a labeling agent, we then examined *in vivo* tracking performance. Approximately twenty radiolabeled B16 cells were introduced into female nude mice (*Foxn1^nu^*) via intravenous or intracardiac injection (Fig. 3). Following injection, mice were imaged using PET (10 min acquisition), and images were reconstructed using the conventional OSEM algorithm provided by the vendor (Sofie Biosciences). After adjusting the display window, we were able to visualize punctate signals originating from various organs within the mice. Consistent with our previous study,^12^ these signals appeared as discrete focal spots (green arrows; Fig. 3a**,c**), closely resembling the imaging pattern observed when imaging *in vitro* samples. The signals were confined to the body of the mice and were clearly distinct from high-frequency reconstruction noise (blue arrows), which was also visible in these images near the edge of the field of view. Typically, reconstruction noise is confined to 1-2 adjacent pixels, whereas the signals observed in these PET images are Gaussian in shape and spread over a larger number of pixels. Additionally, cell signals appear at consistent locations across multiple frames, whereas reconstruction noise are stochastic. Based on these considerations, it is evident that these observed PET signals represent FDG-labeled cells.

**Figure 3.**
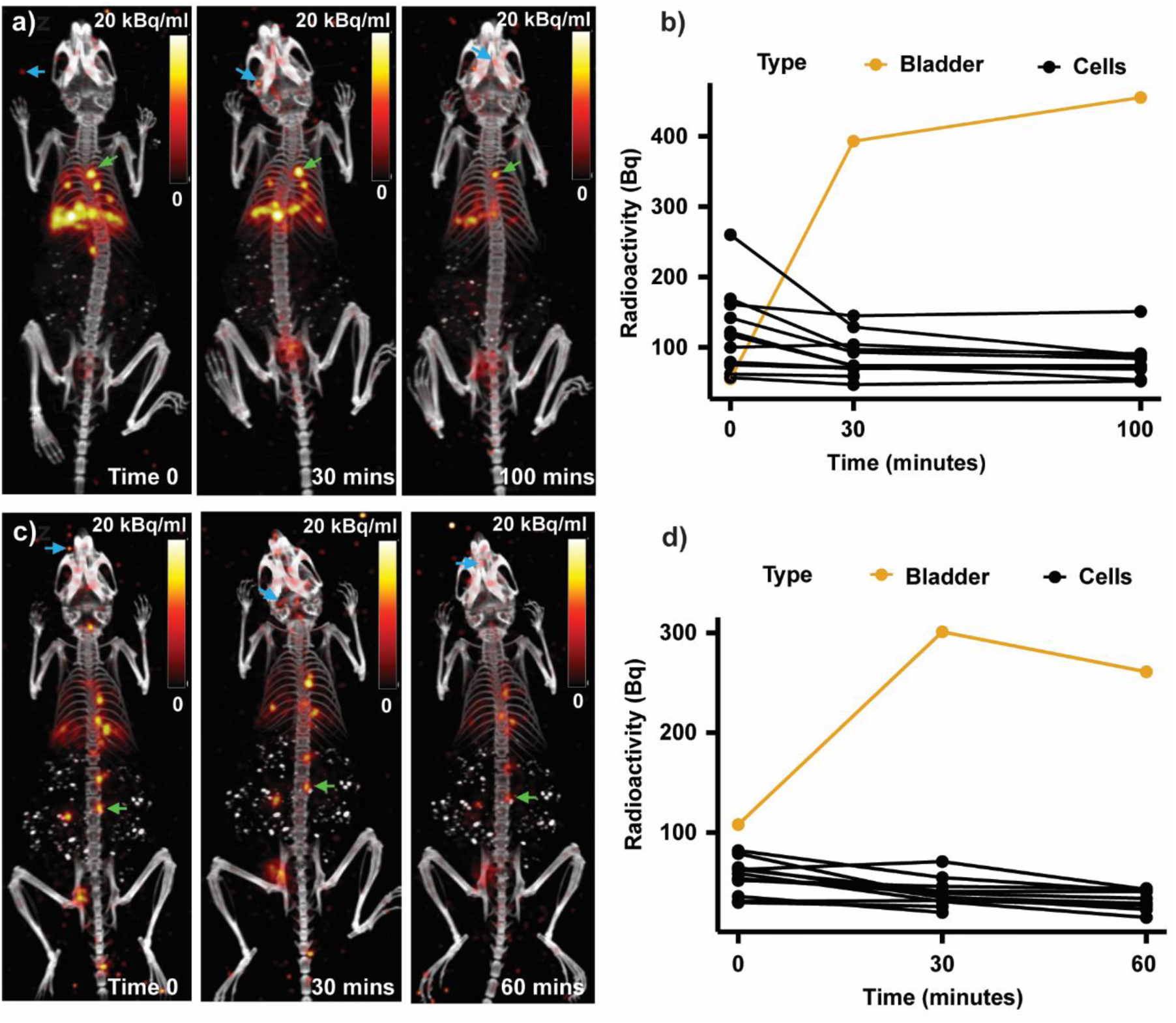
*In vivo* PET imaging of FDG-labeled single cells. **(a, c)** PET/CT images of radiolabeled single cells, which were introduced into mice via intravenous (a) or intracardiac injection (c). PET images were reconstructed using the conventional OSEM method. The focal signals seen in the PET images represent single labeled cells (green arrows). High-frequency reconstruction noise is also visible in the images near the edge of the field of view (blue arrows). **(b, d)** Region-of-interest quantification of the PET images, showing the change in radioactivity in single cells and bladder over time.

As would be expected, dramatically different patterns of cellular arrest arose from different injection routes. After intravenous injection (Fig. 3a), the administered cells were confined to the lungs, which is the first capillary network on their path, whereas after intracardiac injection (Fig. 3c), the cells appeared to spread across multiple organs. It is important to note that about 10-20 cells could be simultaneously visualized within each animal, supporting the feasibility of tracking multiple single cells *in vivo* using PET.

Previously, Jung et al. reported intravenous injection of 100 cells labeled with ^68^Ga-MSNs.^12^ However, in their study, it was difficult to distinguish individual labeled cells from background signal, partly because of the low activity of the cells (average ∼3 Bq/cell). In our study, we could consistently label 100% of B16 cells with >20 Bq. These FDG-labeled cells were remarkably robust, producing a stable signal for up to 100 minutes post-injection. In addition, the PET images had a minimal background in the mouse body, apart from a small amount of FDG visible in the mouse bladder (Fig. 3a**,c**).

To characterize the labeled cells *in vivo* over time, we monitored the radioactivity of single cells by quantifying their radioactivity within regions of interest (ROIs) at different time points (Fig. 3b**,d**). Immediately after intravenous and intracardiac injections, the mean radioactivity per cell was 122 and 56 Bq, respectively. These numbers rapidly dropped to 86 and 38 Bq after 30 minutes, but then seemed to stabilize at around 82 Bq/cell and 33 Bq/cell at later timepoints. This decrease was accompanied by increasing PET signal in the bladder, where activity from cell efflux and a small number of dying cells accumulated over time. From these data, we conclude that conventional OSEM reconstruction can monitor the migration of single cells to various sites throughout the body, using PET data from a 10 min scan.

### Enhanced detection of single cells using the PEPT-EM algorithm

While our results show that OSEM can be used to visualize single cells in vivo, we aimed to further improve the sensitivity of PET for this demanding application by investigating the performance of an algorithm recently developed for tracking multiple moving particles in industrial processes and other opaque systems. Positron emission particle tracking (PEPT) is a method developed on the premise that conventional PET is not optimal for localizing a small number of weakly radioactive point sources.^20–22^ Indeed, PET requires data to be reconstructed into a large tomographic image composed of millions of voxels, which does not efficiently represent point sources. In contrast, PEPT calculates the 3D positions of the sources directly from the coincident annihilation photons recorded by the scanner without explicitly reconstructing an image of the radiotracer distribution. Recently, an expectation-maximization formulation (PEPT-EM) was derived to reconstruct source locations based on a maximum-likelihood approach and could track up to 80 moving simulated particles simultaneously.^23^ However, the method has yet to be investigated for tracking cells *in vivo*.

The difference between PEPT-EM and conventional ordered-subset expectation maximization (OSEM) is illustrated in Fig. 4a**,c**. Both algorithms are iterative, starting with an initial estimate of the image or source positions, followed by several iterations to update the initial guess according to the measured PET data. In the case of OSEM, the input data are binned into sinograms, and the outputs are tomographic PET images. OSEM alternates forward-and back-projections to maximize the agreement between the estimated and measured sinograms. PEPT-EM, on the other hand, operates directly on the raw list-mode data, first soft-clustering the recorded coincidence events according to their proximity to discrete radioactive sources, then updating the 3D positions of these sources based on a distance minimization procedure. Theoretically, two nearly intersecting lines of response are sufficient to determine the location of a radioactive source. However, in practice, many more LORs are required to cope with various sources of uncertainty, including intrinsic background radioactivity and photon scatter.

**Figure 4.**
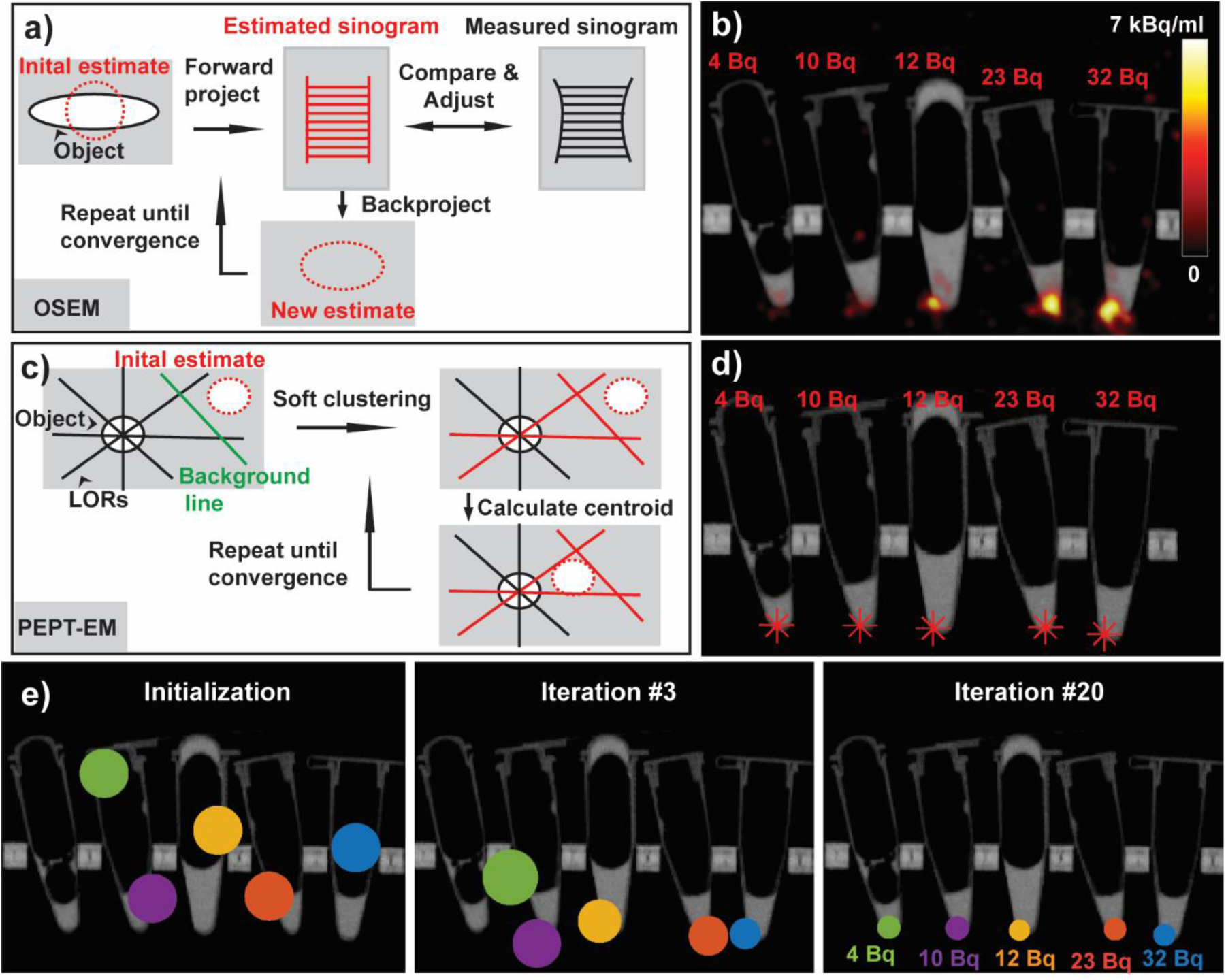
PEPT-EM improves the detection of single cells. **(a)** Schematic representation of OSEM, the standard algorithm for PET reconstruction. **(b)** PET image obtained by OSEM reconstruction, showing five FDG-labeled cells imaged in vials. The lowest detectable cell had 12 Bq. **(c)** Schematic representation of PEPT-EM, a tracking algorithm based on a Gaussian mixture model that estimates the 3D positions of radioactive sources directly from the recorded coincident annihilation photons. **(d)** The 3D positions of the discrete sources were reconstructed by PEPT-EM from the same PET dataset, shown here as red asterisks over the CT image of the vials. **(e)** The PEPT-EM algorithm was initialized by generating random cell locations. As the algorithm iterates, it progressively converges towards the maximum-likelihood location of the cells, leading to a significant reduction in the standard deviation of the estimated position (represented by the radius of the circle).

To evaluate the performance of the PEPT-EM algorithm for tracking single cells, we labeled five B16 cells with FDG, dispensed them in individual vials, then imaged them using PET (Fig. 4b**,d**). Using conventional OSEM reconstruction, we observed focal PET signals for cells containing as little as 12 Bq but could not resolve weaker cells. In comparison, PEPT-EM could correctly localize all five cells, including one containing only 4 Bq. Hence, PEPT-EM is more sensitive than OSEM for localizing weak point sources such as radiolabeled single cells.

To illustrate the convergence of the algorithm towards its solution, we plot the estimated positions of these 5 cells after 0, 3, and 20 iterations (Fig. 4e**; Supplementary Vid. 1**). Here, the center of each circle represents the estimated position of each cell, whereas the radius represents the standard deviation of the estimate. The PEPT-EM algorithm was initialized assuming random cell locations. As the algorithm iterates, it progressively clusters the list-mode data into different groups based on the probability that each detected event originated from each of the different cells being tracked. As the algorithm converged towards the maximum-likelihood location of the cells, the standard deviation decreased from an initial value of 5 mm down to ∼1.0 ± 0.1 mm. The most radioactive cell (32 Bq) converged the fastest and took only 3 iterations to settle, whereas the weakest cell (4 Bq) converged last, after 20 iterations.

We next used PEPT-EM to track the dissemination of melanoma cells *in vivo.* Around 20 FDG-labeled B16 cells were injected into nude mice (*Foxn1^nu^*) via intravenous or intracardiac routes, then PET/CT scans of these mice were acquired in list-mode format. The PET data were first reconstructed using the OSEM method to produce conventional tomographic images showing the distribution of the tracer in 3D. As previously shown, after intravenous injection, melanoma cells could be clearly seen in the lungs (Fig. 5a**, Supplementary Vid. 2**), whereas, after intracardiac injection, they were widely disseminated throughout the entire body and could be observed in many different organs (Fig. 5b**, Supplementary Vid. 3**). We then reconstructed the list-mode data using PEPT-EM to derive the precise 3D single-cell locations for both injection routes.

**Figure 5.**
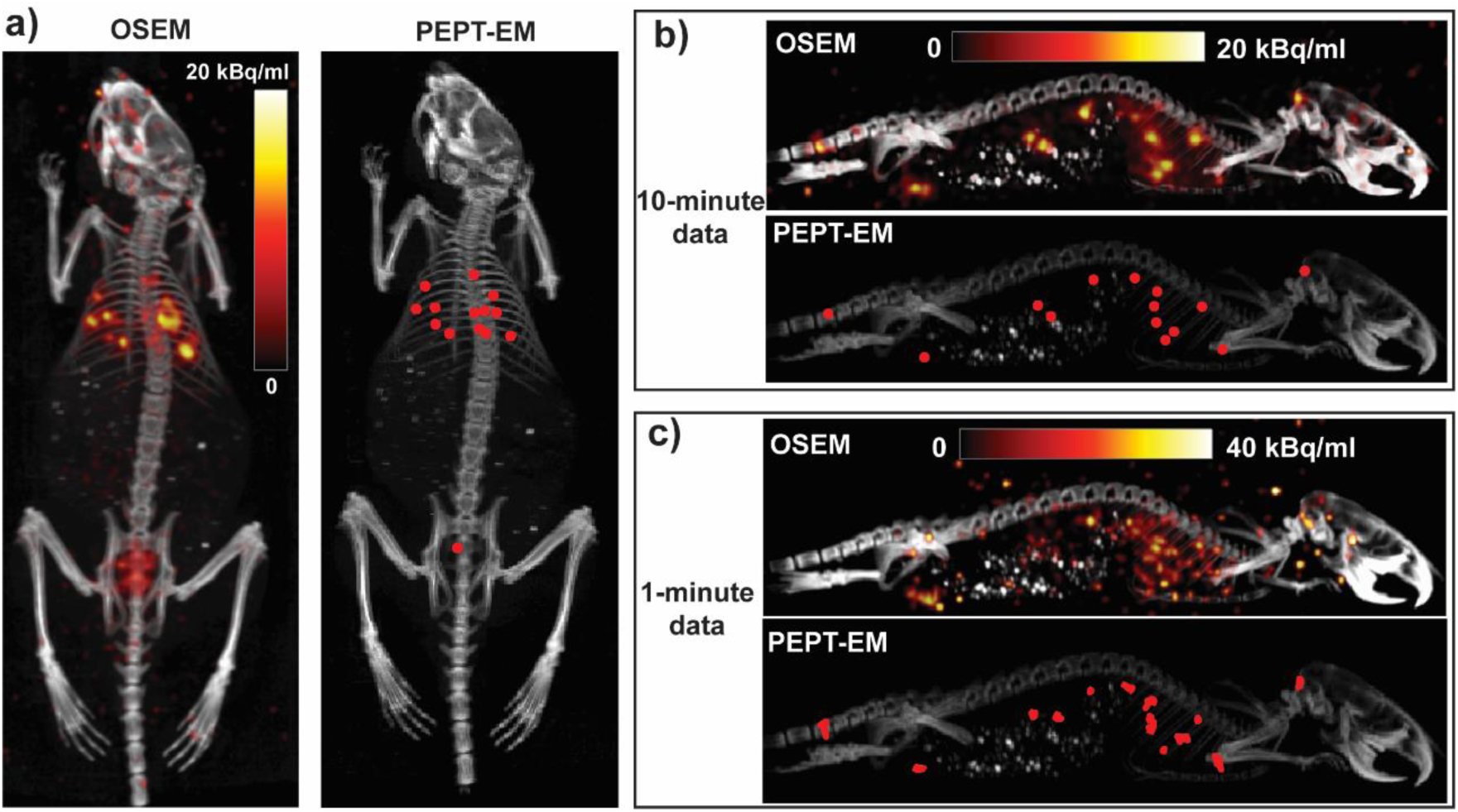
*In vivo* tracking single cells using PEPT-EM. **(a)** The PEPT-EM algorithm was applied to a PET dataset acquired after intravenously injecting ten to twenty single cells into *Foxn1^nu^*mice. The resulting cell positions (red asterisks; right panel) are compared to the conventional OSEM reconstruction of the same dataset (left panel). **(b-c)** Both algorithms were also applied to a PET dataset obtained by injecting FDG-labeled cells into the left ventricle of a mouse. The results are shown for 10-minute (b) and 1-minute acquisitions (c).

Overall, the cell locations estimated by PEPT-EM matched the positions where focal signals were observed in the OSEM-reconstructed PET image (Fig. 5a**,b**). Because of physiological FDG accumulation in the bladder, PEPT-EM automatically assigns one of the sources to this organ. Unlike OSEM, PEPT-EM directly provides an estimate of the number of sources in the field of view and their 3D position. As the exact number of injected cells is not exactly known, we ran PEPT-EM assuming a larger number of cells (90 simulated sources) and filtered out any reconstructed source that did not meet specific requirements in terms of minimum radioactivity and cluster variance. In general, the PEPT-EM algorithm does not converge to a single global solution. However, by randomly initializing the algorithm using more sources than physically present in the animal, we can better sample the solution space and improve the convergence of the algorithm towards a global solution. Here, the initial source positions were randomly generated near the center of the mouse. To confirm that the outcome of the algorithm is independent of its initialization, we reconstructed the intracardiac dataset four times using different random initial positions and confirmed that the same solution is achieved independently of the initialization (**Supplementary** Fig. 7). Moreover, the same solution was obtained regardless of whether the source positions were initialized near the center of the mice or randomized over the entire body. **Supplementary Vid. 4** illustrates the convergence of the algorithm over 300 iterations.

Finally, we analyzed the performance of PEPT-EM under low-count conditions, which we simulated by splitting the 10-minute PET acquisition into ten frames of 1 minute. When applied to the smaller datasets, PEPT-EM yielded 3D cell positions that were consistent with the full 10 min dataset. The computed positions appeared to fluctuate slightly from frame to frame, but the variations were within the spatial resolution of the system and likely caused by statistical noise rather than physiological cell motion (Fig. 5c**, bottom**). In contrast, due to the low statistics, OSEM reconstruction of the 1-minute acquisition was noisy and failed to resolve single cells (Fig. 5c**, top**). This result confirms our previous observation that PEPT-EM is more sensitive for tracking weakly radioactive single cells.

### The fate of radiolabeled single cells following intracardiac injection

Intracardiac injection is commonly used as an experimental mouse model of bone metastasis.^24–26^ However, the procedure is technically challenging, and a high degree of precision is required to inject the cells into the small and pulsing volume of the left cardiac ventricle (**Supplementary Vid. 5)**. Even with ultrasound guidance, it is difficult to confirm whether an injection was performed correctly. Using our imaging technique, we observed three different scenarios following intracardiac injection: i) The cells were correctly injected into the left ventricle of the mouse and distributed throughout the entire body; ii) The cells were inadvertently injected into the nearby right ventricle and trafficked to the lungs, where they arrested; iii) The cells were injected outside of the heart into the pericardial cavity (**Supplementary** Fig. 8). In this context, the ability of PET to track the fate of single cells could help establish more reliable models of bone metastasis by confirming the successful administration of cells.

To further illustrate the unique capabilities of the proposed cell tracking methodology, we traced the fate of 74 FDG-labeled melanoma cells after successful intracardiac injection in healthy mice. We evaluated the spatial distribution of the arrested cells using co-registered CT images to identify the anatomical sites where single cancer cells were seen to arrest (Fig. 6a**, Supplementary** Fig. 9). These sites were then systematically categorized according to conventional organ systems, distinguished by color-coded groupings (Fig. 6b). We observed that a substantial number of cells arrested in large organ systems, such as lung, musculoskeletal, and digestive systems. The results are consistent with the formation of bone metastases that are known to arise in this animal model.^26–28^

**Figure 6.**
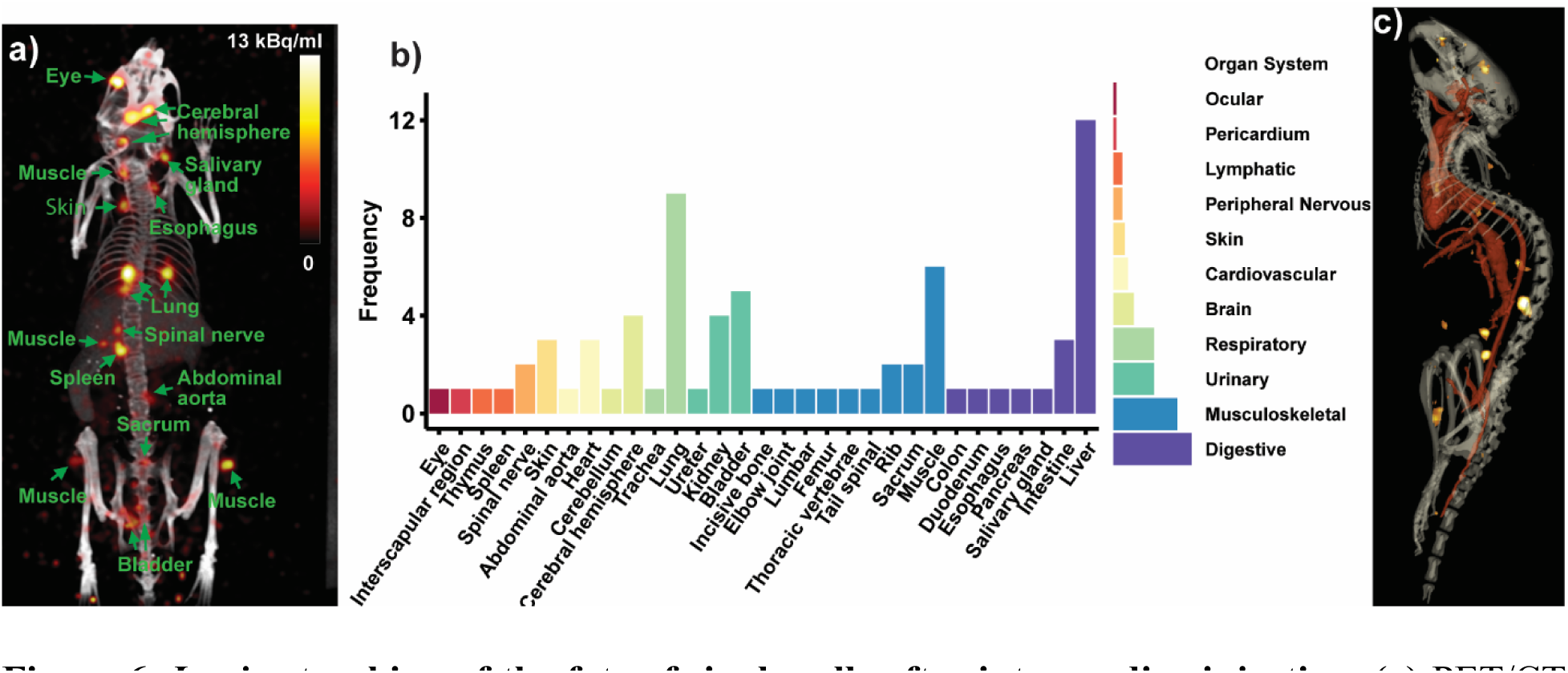
*In vivo* tracking of the fate of single cells after intracardiac injection. **(a)** PET/CT image (OSEM reconstruction; maximum intensity projection) of a mouse right after intracardiac (left ventricle) injection, with labels indicating the anatomical locations where cancer cells were arrested. **(b)** A comprehensive summary of the results, showing the sites and organ systems in which 74 labeled cells arrested (*n*=6 mice). **(c)** Three-dimensional rendering showing single-cell distribution (OSEM reconstruction) relative to the segmented bony and cardiovascular anatomy.

Finally, we examined the pattern of cell arrest in relation to the vascular system, which was imaged using a nanoparticle-based CT contrast agent. The agent, which was introduced intravenously in mice after intracardiac injection of cancer cells, remained detectable in the bloodstream for 30 minutes, with a peak observed at around 15 minutes (**Supplementary** Fig. 10). Due to the 100 µm resolution of this scan, we could clearly resolve larger arteries such as the aorta, carotid arteries, and jugular veins, but not smaller tissue capillaries. However, the location of the arrested cells seen on PET did not correlate with the larger vessels visible on CT. This observation suggests that the labeled cancer cells were likely trapped in the smaller capillaries within the tissues (Fig 6c**, Supplementary Vid. 6**).

## 3. Discussion

Taken together, our results conclusively show that PET is sensitive enough to track the individual fate of 10 or more cells simultaneously in the same animal. This study represents a significant improvement over our previous work,^12^ where we could only track cells one at a time. The improved tracking workflow solves a crucial bottleneck by enabling multiple cells to be tracked simultaneously, dramatically increasing the amount of information that can be garnered through each experiment. This crucial benefit is vividly illustrated in our analysis, which tracked the fate of 74 single cells in 6 mice (Fig. 6). If performed using our prior single-cell method and considering the success rate for intracardiac injection, such a study would have required preparing and imaging well over 100 mice, which would have been prohibitive.

In addition, the workflow reported herein solely relies on commercially available reagents and instruments. Contrary to expectations, metabolically active cancer cells (e.g. B16) could be reliably labeled with 25-100 Bq of FDG per single cell, an amount largely sufficient for high-accuracy tracking (Fig. 2c). The labeling protocol is straightforward and involves only short-term cell fasting and incubation with FDG in glucose-free media, without any nanoparticle radiochemistry. Additionally, ^18^F is advantageous from the perspective of imaging physics as it possesses a shorter positron range (1 mm) than ^68^Ga (2.9 mm), which improves tracking accuaracy.^29^ This new workflow also benefited from the use of a high-sensitivity, high-resolution, low-background, dual-layer (BGO/LYSO) preclinical PET scanner to image the labeled cells,^15,16^ which considerably improved the detectability of weakly labeled cells. Lastly, we used a microfluidic device to select and dispense fixed numbers of FDG-labeled cells, which improved reliability and reduced the preparation time to less than 10 minutes.

In addition to being more straightforward, FDG labeling demonstrated performance comparable to nanoparticle-based labeling. FDG-labeled cells appeared as tiny hot spots, closely resembling the previously observed pattern of cells labeled with ^68^Ga-MSNs. These cells maintained high metabolic activity (>75% for up to 54 hours; Fig. 2e), a figure, however, slightly lower than for cells labeled with ^68^Ga (90% for up to 50 hours). This lower metabolism after FDG labeling is likely caused by DNA damage, which was equivalent to an X-ray dose of over 20 Gy (Fig. 2g**,h**). DNA damage is known to cause transient cell cycle arrest, which could explain the observed decrease in metabolism. However, given the short half-life of ^18^F, tracking experiments are limited to a short time frame of a few hours, during which the effects of radiation exposure have not yet manifested themselves (**Fig. 2d,f, Supplementary Fig. 4**). In the rare instance that a cell does not survive, the ensuing release of FDG from its cytosol would cause its PET signal to vanish from the image (Fig. 2b), thus preventing dead cells from being included into further downstream analysis.

Another major finding is that PET could detect single cells labeled with as little as 4 Bq. This unprecedented level of sensitivity was achieved using a high-performance PET scanner for data acquisition and leveraging the strength of the PEPT-EM algorithm for cell detection. Using conventional OSEM image reconstruction, we achieved a limit of detection for single cells of 12 Bq using a dual-layer BGO/LYSO microPET scanner with 10.4% sensitivity,^15,16^ which compares favorably to the limit of 28 Bq previously achieved using a single-layer LSO PET scanner.^12^ The detection limit was further lowered by reconstructing PET data using the PEPT-EM algorithm, which was developed to localize sources directly from raw list-mode PET data. Unlike conventional OSEM, PEPT-EM accounts for the discrete nature of the radioactive sources being tracked, thus turning an image reconstruction problem into a source localization problem that is better suited to the task of tracking single cells. In turn, the improved detection limit of PEPT-EM translates into higher sensitivity for tracking single cells *in vivo*, which was particularly evident for low-count data (Fig. 5b**,c**). Further lowering the limit of detection would extend the applicability of this technique to a broader range of applications, including tracking cells with low radioactivity or over long periods of time and monitoring radiosensitive immune cells.

While this study focused primarily on imaging static cells, being able to image cells while they move would provide crucial insight into the dynamics of cell trafficking. Our results show that PEPT-EM could simultaneously track up to 14 single cells (mean radioactivity of 56 Bq/cell) with a temporal resolution of 1 minute (Fig. 5c). PEPT-EM treats dynamic data as a series of independent frames, which limits its ability to track the motion of low-activity sources. In their initial demonstration, Manger et al. used PEPT-EM to track the rotation of 10 ion-exchange resin particles labeled with 1-10 MBq radioactivity at velocities of up to 50 cm/s.^23^ Previously, our group developed a custom algorithm, CellGPS, for tracking the dynamics of a single cell *in vivo*.^12,13^ Using the CellGPS algorithm, we tracked the motion of a single cell (70 Bq) traveling at a maximum velocity of 5 cm/s, from its injection site in the tail vein to its ultimate arrest within pulmonary capillaries. Unlike PEPT-EM, the CellGPS algorithm models the trajectory of a moving cell as a continuous function of time and can thus reconstruct complex spatiotemporal trajectories from fewer detected counts. However, CellGPS is unable to track more than one cell at a time. We are currently investigating the feasibility of combining PEPT-EM and CellGPS into a single algorithm to improve the temporal resolution for multiplexed cell tracking.

To the best of our knowledge, this study is the first to examine the distribution of individual cancer cells throughout the entire body after intracardiac injection. Through this improved method, we could localize disseminated melanoma cells throughout the body with excellent spatial resolution.

From co-registered CT images, we identified the organs where migrating cells tended to arrest and found that B16 cells were widely distributed throughout the mice, reaching 31 organs and 11 organ systems. This finding was expected given that the left cardiac ventricle supplies arterial blood to the entire body. Our data further suggest that cells arrested very quickly after injection, within the first minute, likely through passive trapping in capillary beds. Interestingly, prior studies of metastases formation following intracardiac injection mainly reported bone metastases,^24,27,28^ even though the musculoskeletal system accounted for only 22% of the cell arrest in our study. This observation suggests that the bone marrow may provide a more supportive niche for the survival and rapid growth of metastatic lesions, as originally postulated in the “seed and soil” hypothesis.^30,31^

While several advantages of FDG-PET for tracking single cells have been described, a few limitations should also be mentioned. The short decay half-life of FDG of 110 minutes limits the tracking time frame to a few hours and hence prevents FDG from being used for tracking cells over multiple days, including, for instance, for tracking the migration of hematopoietic stem cells during bone marrow transplant or immune cell homing in response to inflammatory stimuli.^4,32^ FDG is better suited for short-term tracking applications, as reported in this study. For long-term tracking of radiolabeled cells, long-lived isotopes such as ^89^Zr (3.3 days half-life) are available to track cells over 7 days.^12^ FDG may also be less efficient as a label for cells with low basal glucose metabolism, such as quiescent immune cells and stem cells. A suitable radiolabel should be selected considering the characteristics of the cells being tracked. Another limitation of FDG labeling is that cells suffered DNA damage from exposure to radiation during the 1 h incubation with the radiotracer (370 MBq/ml). An alternative labeling method using a microfluidic device could label cells instantly, which would reduce DNA damage by avoiding prolonged radiation exposure during the labeling process.^33–35^

## 4. Conclusion

Tracking single cells within living subjects is crucial to better understand the dynamic trafficking patterns that underlie key biological phenomena such as metastasis and immune cell homing. This is the first demonstration that PET can track 10-20 single cells simultaneously, a finding that holds significance for cell-tracking applications in biological research and cell-based therapies. This achievement was made possible through improvements in cell labeling, cell isolation, cell imaging, and data reconstruction. To facilitate its routine use and dissemination, the workflow was implemented using commercially available products and reagents. Furthermore, we demonstrated that PET could detect single cells labeled with as little as 4 Bq. As a result, tracking of FDG-labeled single cells *in vivo* could be achieved for low-count data (1-minute acquisition time), which was not attainable using conventional OSEM image reconstruction. Finally, as a proof of concept, we investigated the fate of melanoma cells following intracardiac injection into mice. Most cells accumulated in large organ systems, such as the pulmonary, musculoskeletal, and digestive systems, and away from primary large vessels. Overall, this study highlights the potential of PET imaging for whole-body cell tracking applications during disease progression and cell-based therapies at the single-cell level, offering unmatched sensitivity and unique insight into the earliest phases of cell trafficking.

### Methods Cell Culture

Two murine cancer cell lines (B16-F10 and 4T1) and one human cancer cell line (MDA-MB-231) were originally acquired from American Type Culture Collection (ATCC). We cultured B16-F10 and MDA-MB-231 in Dulbecco’s Modified Eagle Medium (DMEM) medium (ThermoFisher Scientific, #11995065), and 4T1 cells in Roswell Park Memorial Institute (RPMI)-1640 medium (ATCC, #302001). Unless otherwise noted, both cultured media were supplemented with 10% fetal bovine serum (FBS) (ThermoFisher Scientific, #26140079) and 1% penicillin-streptomycin (ThermoFisher Scientific, #15140122). The cells were cultivated in T25 culture flasks inside a humidified incubator at 37°C and 5% CO_2_ until confluence and transferred to tissue culture plates for radiolabeling and biological assays. Cells used for radiolabeling and biological assays did not exceed passage 15.

### Radiolabeling with FDG

B16-F10, 4T1, and MDA-MB-231 were first seeded into a 6-well plate at 2×10^5^ cells/well and were cultured for 48 hours. To enhance FDG uptake, these cells were kept for 1 h in a glucose-free medium (DMEM medium with L-Glutamine; ThermoFisher Scientific, #11966025) and 1% FBS prior to radiolabeling. After this fasting period, we replaced the fasting media with fresh fasting media mixed with FDG (370 MBq/ml, 2 ml per well). Clinical-grade FDG was produced at the Stanford Cyclotron and Radiochemistry Facility through nucleophilic ^18^F-fluorination and hydrolysis of mannose triflate. The radiotracer was picked up within 2 hours of production to ensure high specific activity. We incubated cells with FDG for 1 hour, then washed each well three times with 2 ml of PBS to remove residual extracellular FDG.

### Single Cell Dispensing

We used a microfluidic-based single-cell sorter (Hana, Namocell) to dispense single cells into a 96-well plate. We first stained the cells with Calcein-AM (5 nM; ThermoFisher Scientific, #65085381; 8 min incubation) and diluted the cell suspension to achieve 5,000 – 10,000 cells/ml. As per the device’s instruction for dispensing single cells, we loaded the diluted cell solution into a cell cartridge and ran the device in analysis mode to differentiate live cells from dead cells and debris. After analyzing around 200 cells, we gated out the debris population (defined by low forward scattering and side scattering values) and selected viable cells with strong green fluorescence (typically, the threshold was set at 50). We then ran the device in dispensing mode to dispense 30 single cells into the media-prefilled wells of a 96-well plate. Each single cell was dispensed inside a 1 μl droplet by the device.

### Single-Cell Gamma Counting

An automatic gamma counter system (Hidex) was utilized to measure the radioactivity of individual cells. Single radiolabeled cells were transferred to 250 μl microcentrifuge tubes and loaded into racks for automated gamma counting. Each tube contained 100 μl fresh media and 1 μl droplet from the single-cell dispenser (composed of 99.7% PBS sheath fluid, 0.3% cell culture media solution). As a result, the radioactivity in each tube mainly originated from the single cell as the efflux in the cell solution was highly diluted. The radioactivity values in counts per second were converted to Bq through a calibration curve constructed from the measurement of serially diluted FDG solutions and a standard dose calibrator (Biodex, Atomlab 400). To confirm the singularity of dispensed single cells, we divided the solution in each tube equally into two or three vials and measured the activity of each fraction by gamma counting and PET imaging. The rationale for this experiment is that the contents of the single cell cannot be fractionated into multiple vials.

### Cell Viability Assays

The effect of radiolabeling on cell metabolism was assessed using the colorimetric Cell Counting Kit 8 (CCK-8) assay. B16 cells were first seeded onto a 96-well plate at a cell density of 5,000 cells per well and grown for 24 hours. FDG labeling of these cells was carried out as described in the previous section, including cell fasting and incubation with FDG for 1 hour (370 MBq/ml, 200 μl per well). We added 10 μl of CCK-8 reagent (Sigma Aldrich, #96992) to each well at times 0, 6, 24, and 54 hours after radiolabeling. Cells incubated with methanol (MeOH) for 10 minutes were prepared as positive control samples. We measured the absorbance of the solution at 450 nm using a plate reader (GloMax multi-detection system, Promega). The metabolism of radiolabeled cells was compared to control cells that were cultured similarly to the labeled cells in all aspects but were not labeled with FDG.

We also used the Sytox Green dye to assess the integrity of the cell membrane immediately after radiolabeling. While performing the radiolabeling procedure mentioned in the previous section, we added 2 μl of Sytox Green dye (ThermoFisher Scientific, NucGreen #R37109) to 500 μl of FDG-labeled B16 cell suspension for 15 minutes at room temperature. The excess dye was removed, and the Sytox-stained cells were quantified by flow cytometry using the single-cell sorter (Hana, Namocell). B16 cells fixed using ice-cold Ethanol for 15 minutes were prepared as the positive control for this assay.

### Apoptosis Assay (Annexin-V staining)

Annexin-V was utilized to measure apoptosis after radiolabeling. B16 cells were seeded into a 6-well plate (2×10^5^ cells/well) and cultured for 48 hours. The cells were then labeled with FDG as previously described. Excess FDG was removed, and the radiolabeled cells were stained with an Annexin-V staining kit (Abcam, #ab176749) three hours after radiolabeling. Annexin-V labeled cells were quantified by flow cytometry using a single-cell sorter (Hana, Namocell). B16 cells treated with Cisplatin (100 μM; Sigma Aldrich, #P4394) for 48 hours were prepared as a positive control for this assay.

### DNA Damage Assay (γH2AX staining)

We used γH2AX immunostaining to characterize DNA damage caused by radiolabeling. We first seeded 1.5×10^5^ B16 cells into glass bottom dishes (iBidi, #81218-200) and cultured them for 36 hours. Following the FDG labeling procedure, we stained FDG-labeled and control cells against γH2AX at 0 and 24 hours post-labeling. The immunostaining procedure started with fixing the cells in methanol on ice for 5 minutes, followed by cell permeabilization with 0.1% Triton-X 100 in PBS for 10 minutes and blocking in 1% BSA / 0.1% Tween 20 in PBS for 30 minutes. The cells were then stained using a primary antibody against γH2AX (1:100 dilution; Sigma-Aldrich, #05-636) in the blocking solution at 4°C overnight. Secondary staining was carried out using an anti-mouse Alexa Fluor 488 antibody (1:100 dilution; ThermoFisher Scientific, #A-21202) in PBS. The cell nuclei were stained with DAPI (1:50 dilution; ThermoFisher Scientific, NucBlue #R37606) in PBS. As an additional reference, we irradiated cells (10 and 20 Gy) using X-ray (225 kV; 13 mA; X-RAD SmART irradiator, Precision X-Ray Inc). These irradiated cells were stained 10 minutes following radiation, together with control unirradiated and labeled cells. Finally, the stained cells were imaged using a fluorescence microscope (EVOS FL, 40x objective in oil), and mean fluorescence intensity was quantified using ImageJ software.

### Cell Migration Assay

We used the wound healing assay to examine the ability of radiolabeled cells to proliferate and migrate in response to a mechanical scratch of the cell monolayer. We first seeded B16 cells into a 6-well plate at 1.5×10^5^ cells/well and cultured the cells for 36 hours. B16 cells were radiolabeled with FDG as described in the previous section, then a gap of 0.8-1 mm width was created by scratching the cell monolayer with a narrow cell scratcher. The process of closing the gap was observed with a brightfield microscope (EVOS FL, 4x objective) at 0, 6, 24, and 48 hours after gap creation. B16 cells treated with Cisplatin (100 μM; Sigma Aldrich, #P4394) for 48 hours were prepared as the positive control for this assay.

### Radiolabeling Efflux

To examine the FDG efflux following radiolabeling, we seeded B16 cells onto a 96-well plate at a cell density of 15,000 cells per well and grew the cells for 24 hours. FDG labeling was performed per our standard protocol (370 MBq/ml, 200 μl per well). We removed excess FDG and washed the cells thrice with PBS (100 μl each time). We collected the supernatant and the lysed cells at times 0, 30, 60, 90, and 120 minutes after the removal of FDG. The radioactivity of these samples was measured with a gamma counter (Hidex). We characterized the efflux of FDG in three different culture media with different concentrations of D-glucose (0, 1.5, and 4.5 g/L). The FDG retention percentage was computed as the ratio of the radioactivity in lysed cells over that in both lysed cells and supernatant.

### Injection of radiolabeled single cells *in vivo*

All *in vivo* procedures in this study followed the protocol approved by the Administrative Panel on Laboratory Animal Care at Stanford University. We injected FDG-labeled single cells into athymic nude mice (*Foxn1^nu^*, female, 6-8 weeks old, strain# 490, Charles River) via tail vein (intravenous) and left cardiac ventricle (intracardiac) routes. The mice were anesthetized using 2% isoflurane concentration and 0.5 L/min oxygen flow rate during the injection. The injection volume was 100 – 150 μl and contained around 40 FDG-labeled cells. We estimate that around half of the cells were successfully injected, whereas the other half was lost during transfer due to the dead volume of the syringe and adhesion to the microcentrifuge tube and micropipette tip. For intravenous injection, we first warmed the mouse tail with lukewarm water for 2-5 minutes until the tail vein became visible, then injected the cells using an insulin syringe (BD) attached to a 28g needle. For intracardiac injection, we used an ultrasound system (Vevo 2100, Visual Sonics) to guide the needle to the mouse’s left ventricle in real time.^36^

### PET/CT imaging of radiolabeled single cells

We used a dual-layer BGO/LYSO PET/CT scanner (GNEXT, Sofie Biosciences) to image FDG-labeled single cells in vials and *in vivo*. Mice were anesthetized using 2% isoflurane and 0.5 L/min oxygen flow rate during imaging. PET images were acquired with an acquisition time of 10 minutes and the standard energy window (350 - 650 keV). CT images were acquired using the standard settings with a 2-minute scan time and 80 kVp beam energy. The PET images were reconstructed using the conventional ordered-subset expectation maximization (OSEM) method provided by the vendor with the scanner. We used the OsiriX software to view PET and CT images. Image display was achieved via maximum intensity projection of the PET data using a slice thickness of 6 mm for small vials and 21-22 mm for mice, which allowed us to display all the single cells in a single slice. To quantify the radioactivity of the single cells, we used the Inveon Research Workspace (IRW) software to define spherical regions of interest (ROIs) surrounding the cells and to calculate the activity per cell (obtained by multiplying ROI mean intensity by ROI volume).

### PEPT-EM tracking algorithm

The PEPT-EM tracking algorithm was originally reported by Manger et al. to track the 3D positions of chemical particles labeled with large amounts of radioactivity (> 1 MBq).^23^ Briefly, the PEPT-EM algorithm uses a Gaussian-mixture model and maximum-likelihood expectation maximization to cluster the recorded photon coincidence events according to the estimated position of the cell from which they were most likely emitted. These coincidence events are recorded in list-mode format and are assigned a timestamp and a line of response (LOR, the line formed by two annihilation photons after positron combines with electron). The algorithm is iterative and alternates between clustering the list-mode data and estimating the new positions of the labeled cells until converging to the maximum-likelihood positions of these cells. Additional details of the PEPT-EM algorithm are provided in the supplementary method. The algorithm was implemented using MATLAB (version 2022).

To provide input to the PEPT-EM algorithm, we first had to convert the list-mode data from the machine-generated format to a format readable by MATLAB, validate this conversion method (**Supplementary** Fig. 11), optimize the annihilation photon penetration depth (**Supplementary** Fig. 12), and co-register the 3D positions produced by our algorithm with the reference frame of the GNEXT PET/CT (**Supplementary** Fig. 13). Additional details are provided in the supplementary method. Additionally, the algorithm must be initialized by generating random cell positions. For experiments involving cells in vials, the initial positions were randomly generated within 25 mm of the vial location as seen on CT (**Supplementary Vid. 1**). For *in vivo* experiments, the initial positions were randomized within the central region of each mouse (**Supplementary Vid. 4**). The stopping criteria was set at a maximum of 300 iterations or a total change in the positions of the sources between consecutive iterations of 0.01 mm or less.

### Injection, imaging and segmentation of CT contrast agent

To enhance the visualization of the vasculature in the mouse model, we intravenously injected a CT contrast agent (ExiTron nano 12000; Miltenyi Biotec) into some of the mice. We used the highest resolution settings on the Sofie GNEXT (80 kVp; bin 1; 100 μm voxels) to acquire CT images of the mice 15 minutes after the injection of the CT contrast agent. The CT images were then fused with the matching PET images and displayed using 3D Slicer software. Segmentation of the cardiovascular and bony anatomy was done using ITK-SNAP, which is a semi-automatic threshold-based toolkit.^37^ Most segmentation algorithms require initialization points, followed by setting a singular value to begin the threshold-based region growth. We opted to use Otsu’s multi-threshold method to select a specific value based on the current region of interest. Then, the ITK-SNAP region growth algorithm was used to grow the segment from the previous segment. For the vessels, the threshold was approximately 147-325 Hounsfield units, while for the bone, a lower threshold of 450-670 Hounsfield units was utilized. Lastly, we also conducted a short longitudinal experiment to observe the kinetics of the contrast agent accumulation in the vasculature and various other organs at 15, 30, and 24 hours post-injection.

### Locating single cells after intracardiac injection

We examined 6 mice that were successfully administered with FDG-labeled cells via intracardiac injection (successful rate ∼ 30%). In these mice, we could visualize radiolabeled cells spreading to multiple organs throughout the body. The number of injected cells in each mouse varied from 5 to 40 cells, with a total of 74 cells that could be detected in 6 mice. The anatomical locations of the arrested cells within the body were determined by a board-certified orthopedic surgeon (coauthor YW) by inspecting the PET/CT data using the OsiriX software.

### Statistical Analysis

All analyses comparing differences between groups in this study were conducted with R (Rstudio 2018) via Analysis of Variance (ANOVA) combined with the Tukey test as a post hoc analysis. The notions of significant differences are as follows: * (p < 0.05), ** (p < 0.01), *** (p < 0.001), **** (p < 0.0001).

## Data Availability

### Code Availability

The MATLAB code used to track single cells via the PEPT-EM algorithm and the code to import raw list-mode PET data into MATLAB from the will be uploaded and made publicly available on the first author’s GitHub account at https://github.com/hieung123

## ASSOCIATED CONTENT

### Supporting Information

Supplement materials include supplement methods and supplement figures/videos shared in separate documents.

## ACKNOWLEDGMENTS

This work was supported by the National Institutes of Health (NIBIB R01EB030367). HTMN gratefully acknowledges funding through the Stanford School of Medicine Dean’s Postdoctoral Fellowship, the Stanford Radiation Oncology and Medical Physics Trainee Seed Grant, and the NIH Stanford Molecular Imaging Scholars (SMIS) Program (NCI T32CA118681). The authors sincerely appreciate the fruitful discussion with Stanford Professor Hong Song on intracardiac injection and Dr. Nam Vu (Sofie company) on converting raw data from the GNext PET scanner to LORs data. We would like to thank Stanford SCi^3^ Directors Dr. Frezghi Habte and Dr. Laura Jean Pisani for providing support with molecular imaging equipment used in this study. We are truly thankful to the Stanford Cyclotron and Radiochemistry Facility Director, the late Dr. Bin Shen, technicians Francis Balmaceda and Ka Ho for help with FDG orders, and Namocell employees (Dr. Junyu Lin and Dr. Thompson Lu) for assisting with single-cell dispensing.

## AUTHOR CONTRIBUTIONS

GP contributed to the conceptualization, experiment design, resources, and editing of the manuscript; CKFC contributed resources; ER contributed resources; ND contributed to experiment (intracardiac injections); YW contributed to analysis (single cells distribution after intracardiac injection); CR and BM contributed to analysis (CT segmentation); HTMN contributed to conceptualization, experiment design, experiment, analysis, and manuscript writing.

## Notes

### Competing Interest Statement

The authors have declared no competing interest.

### Summary of Updates

The introduction and abstract were revised.

