## Supplementary Information for "Efficient and multiplexed tracking of single cells using whole-body PET/CT"

**CONTENTS**

**Supplementary Fig. 1 |** PET imaging of FDG-labeled single cells in vials

**Supplementary Fig. 2 |** Validation of the single cell isolation procedure

**Supplementary Fig. 3 |** Radioactivity quantification of single cells using PET image and comparison to standard gamma counting

**Supplementary Fig. 4 |** Characterization of FDG uptake *in vitro* at different glucose levels

**Supplementary Fig. 5 |** Viability assay by Sytox staining of FDG-labeled cells

**Supplementary Fig. 6 |** Wound healing capability (migration assay) of FDG-labeled cells

**Supplementary Fig. 7 |** Single-cell localization with PEPT-EM with three different initializations

**Supplementary Fig. 8 |** Fate of cells after intracardiac injection

**Supplementary Fig. 9|** Single-cell distribution in 5 mice following intracardiac injections

**Supplementary Fig. 10 |** Contrast-enhanced CT imaging of mouse vasculature

**Supplementary Fig. 11 |** Validation of list-mode import tool

**Supplementary Fig. 12 |** Depth of interaction optimization within scintillator crystal

**Supplementary Fig. 13 |** Spatial co-registration of converted list-mode data with vendor-provided PET-CT coordinates

**Supplementary Methods**

**Supplementary Video Captions**

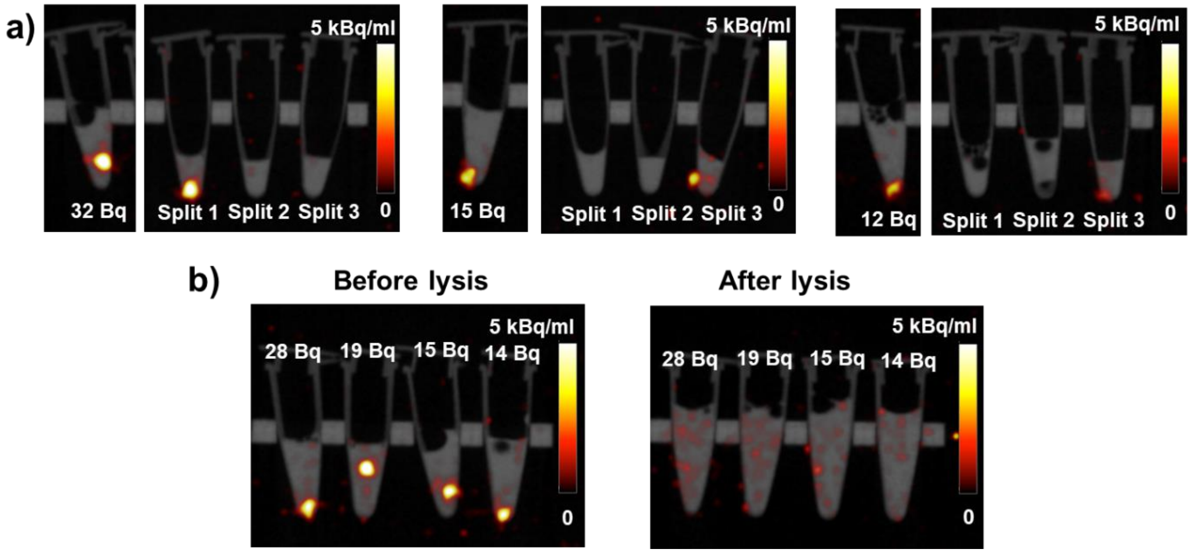


**Supplementary Fig. 1. PET imaging of FDG-labeled single cells in vials. (a)** PET/CT images of three FDG-labeled single cells (activity 12-32 Bq) before and after partitioning into three vials. After partitioning, the focal hot spot was only observed in one of the vials, indicating that the initial focal contrast was due to a single cell. **(b)** PET/CT images of four FDG-labeled single cells (activity 14-28 Bq) before and after adding lysis buffer. The FDG-labeled live single cells appeared as punctate signals, while lysed cells showed diffuse signals from the entire vial. PET images were visualized with OsiriX software as maximum intensity projections, with a slice thickness of 6 mm to capture the cells within the entire vial.

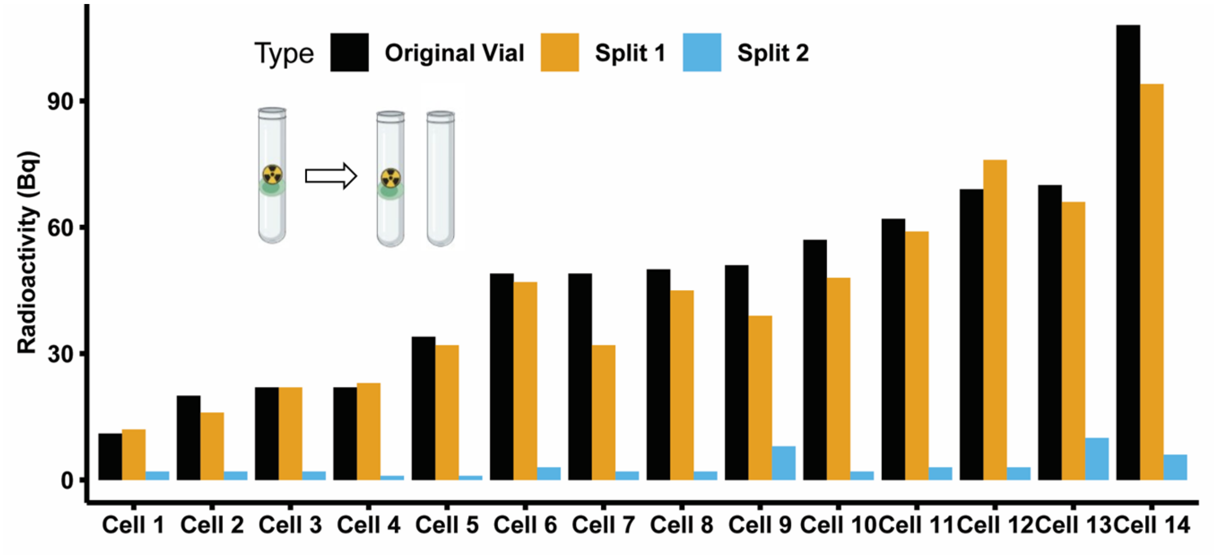


**Supplementary Fig. 2. Validation of the single cell isolation procedure.** Radioactivity of fourteen FDG-labeled single cells (activity 11-108 Bq), measured with a gamma counter before and after splitting into two vials. These data cells were collected over three different labeling experiments, with 4-5 samples in each experiment. Here, we observed a substantial difference in radioactivity values between the two vials after splitting, indicating the presence of a single cell in the original vial. The background radioactivity can be assessed from the measurement of the second vial and is significantly lower than the radioactivity of the single cell.

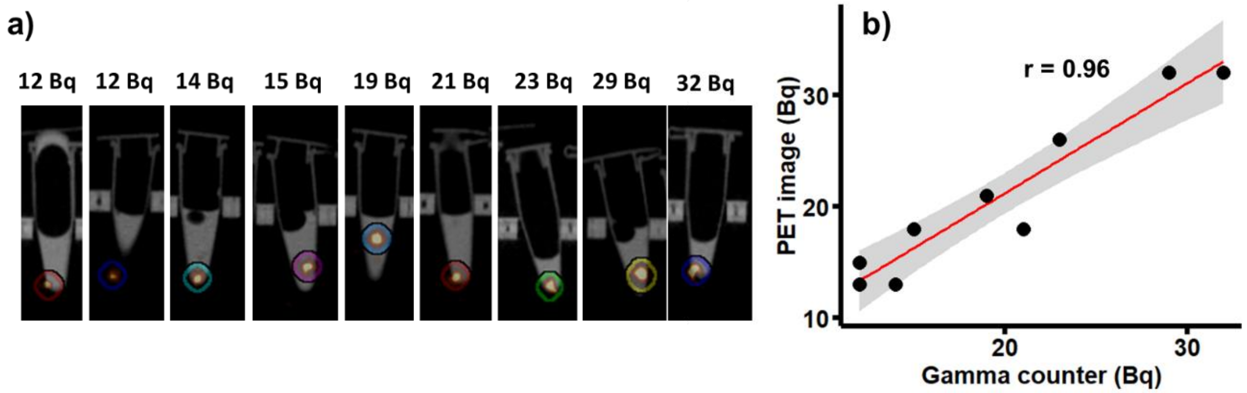


**Supplementary Fig. 3. Radioactivity quantification of single cells using PET image and comparison to standard gamma counting. (a)** PET/CT images of nine FDG-labeled B16 single cells with activities ranging from 12 to 32 Bq (measured by gamma counter). The images were visualized with the Inveon Research Workspace software, and ROI spheres (radius of 5 mm) were drawn around each focal spot to characterize the radioactivity of these cells. **(b)** Scatter plot comparing radioactivity of single cells as measured by PET imaging and gamma counting. The Pearson correlation coefficient between the values estimated by the two methods showed a high degree of positive correlation ($r$ = 0.96). The gray shaded area indicates the 95% confidence interval for linear regression.

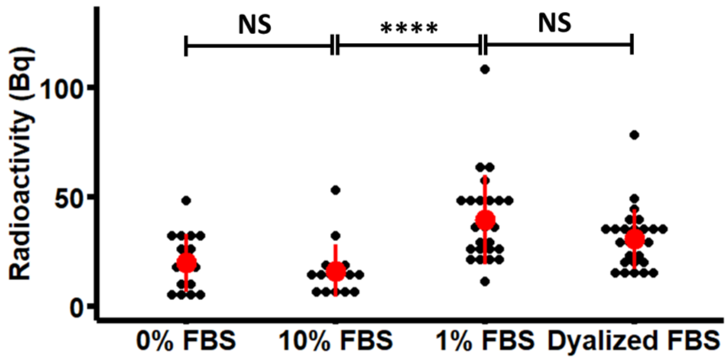


**Supplementary Fig. 4. Characterization of FDG uptake *in vitro* at different glucose levels.** FDG uptake of single B16 cells, measured with a gamma counter, after incubation in media containing 0%, 10%, 1% of qualified FBS, and 10% dialyzed FBS. These concentrations are equivalent to 0, 12.5, 1.25, and 0.1 mg/dL glucose. Glucose influenced FDG uptake even when present in low concentration in FBS. A significant increase was observed when FDG uptake occurred in 1% FBS media (1.25mg/dL glucose) compared to 10% FBS media (12.5 mg/dL glucose). Further reducing glucose concentration using 10% dialyzed FBS (0.1 mg/dL glucose) did not further enhance FDG uptake.

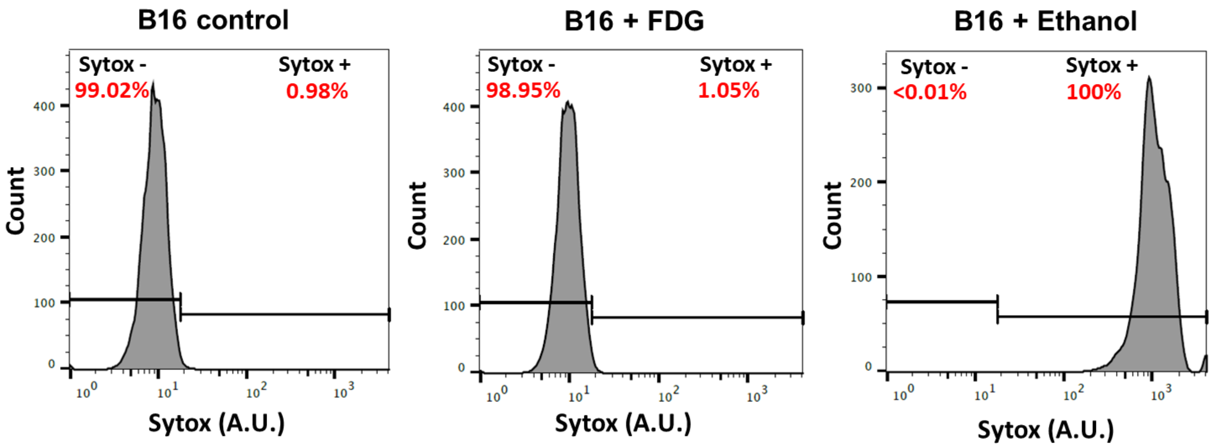


**Supplementary Fig. 5. Viability assay by Sytox staining of FDG-labeled cells.** Three groups of cells (negative control - B16 cells without treatment; FDG-labeled B16 cells 5 minutes after labeling; and positive control - B16 cells fixed with ethanol for 15 minutes) were stained with Sytox (Nucgreen) for 15 minutes at room temperature. We then performed flow cytometry to count the cells positively stained with Sytox signal (indicating cell membrane damage). FDG-labeled cells had similar viability to the negative control group, whereas cells fixed with ethanol were 100% positive for Sytox. These results were repeated in three independent experiments.

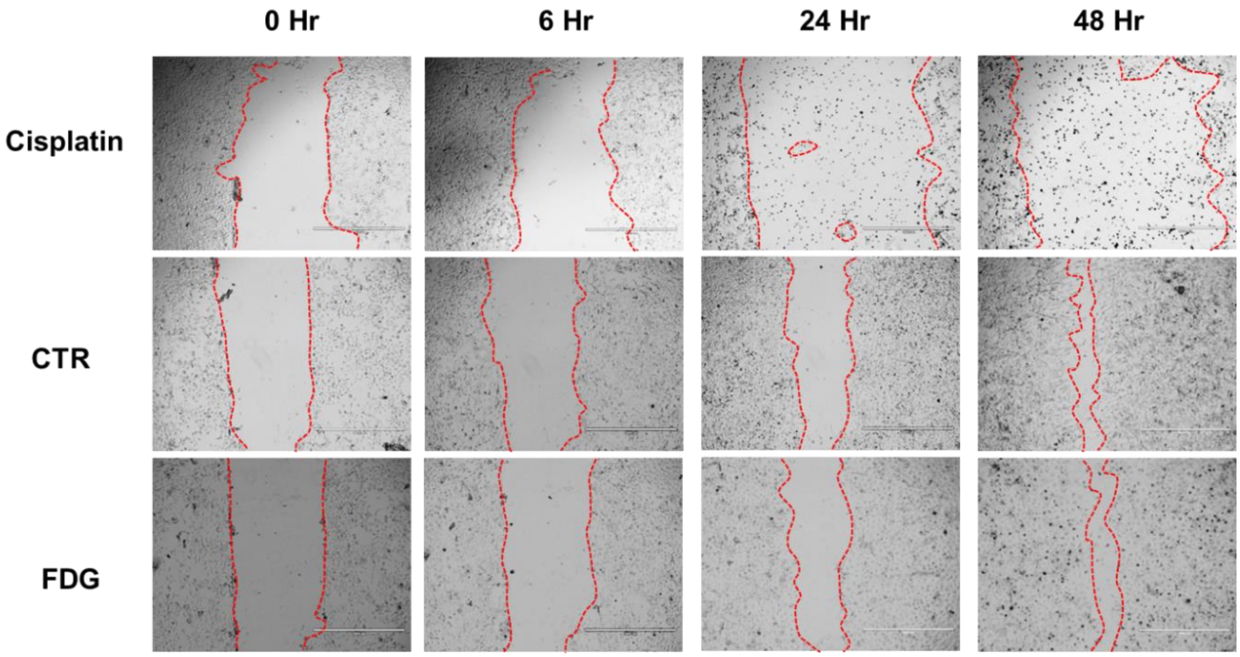


**Supplementary Fig. 6. Wound healing capability (migration assay) of FDG-labeled cells.** Three groups of cells (negative control - B16 cells without treatment; FDG-labeled B16 cells; and positive control - B16 cells treated with Cisplatin 100 μM for 48 hours) were cultured in a 6-well plate. At the initial time point, the plate was scratched to create a small gap (~0.8 mm). We then monitored the migration of cells toward closing this gap at 6, 24, and 48 hours. FDG-labeled cells closed the gap at a rate similar to control cells, while cells treated with cisplatin failed to do so within 48 hours, and in fact, the gap widened. The dots within the cell gap for the Cisplatin sample at 24 and 48 hours were mostly cell debris.

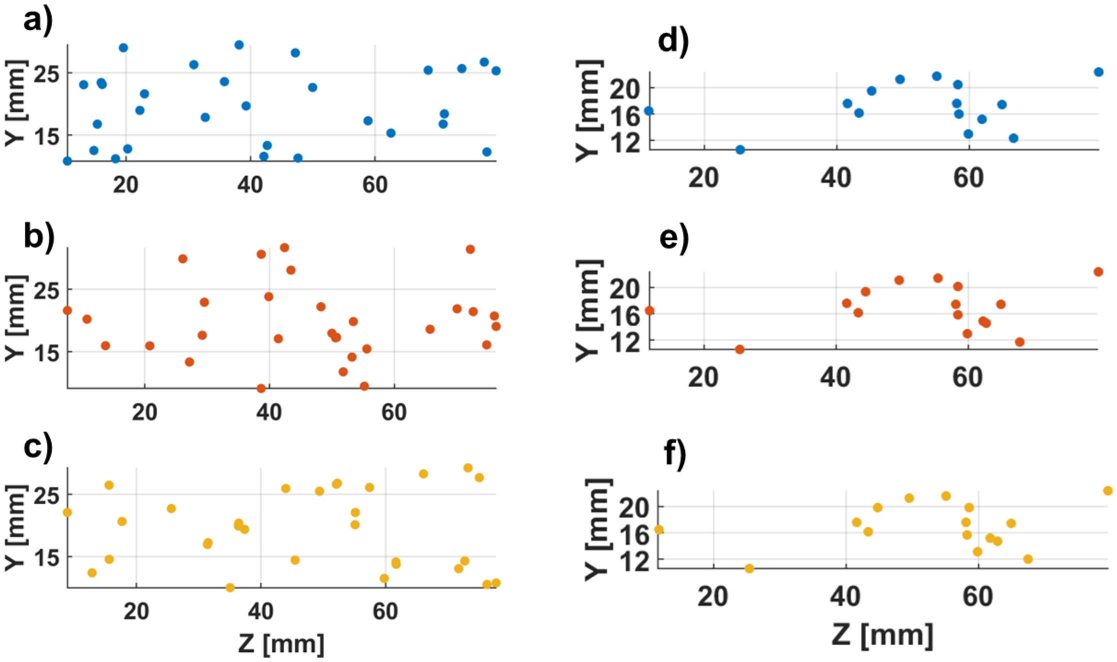


**Supplementary Fig. 7. Single-cell localization using PEPT-EM with three different initializations.** (a-c) Scatter plots of three sets of random initial source positions placed throughout the mouse body. (d-f) Corresponding reconstructed positions obtained using PEPT-EM, for a single-cell PET data set obtained following intracardiac injection. After initializing the tracking algorithm with 30 sources, we filtered out the sources with low activity or high variance at convergence, resulting in 14 single cells distributed throughout the mouse body. The cell distribution patterns were consistent across the three initialization and closely resembled the data reported in Fig. 5b.

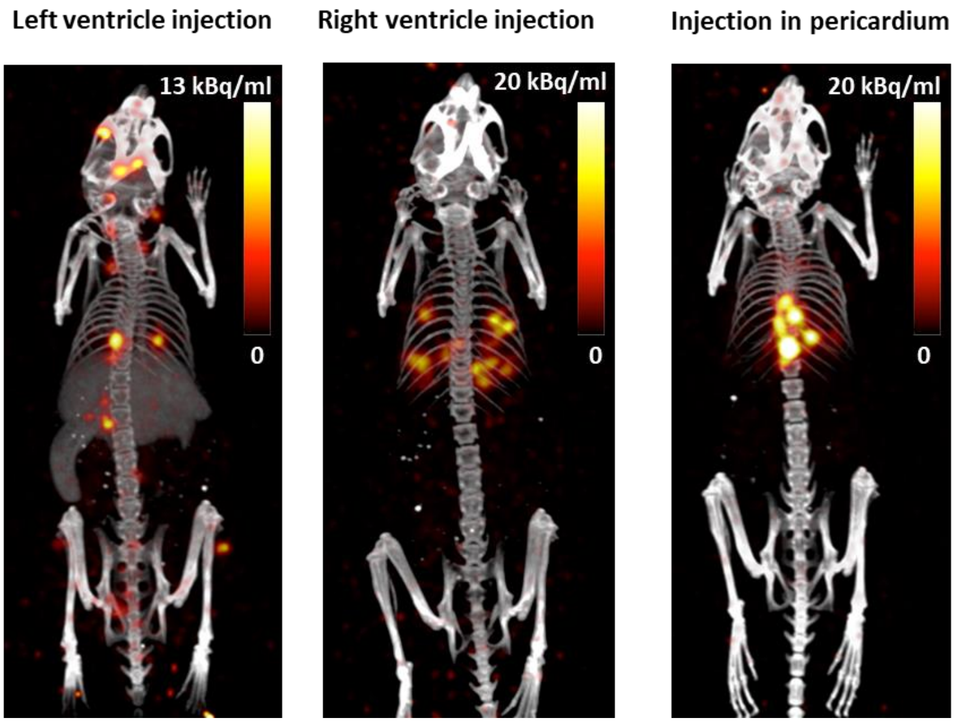


**Supplementary Fig. 8. Fate of cells after intracardiac injection.** PET/CT images illustrating the three possible outcomes following intracardiac injection of 10-20 FDG-labeled B16 cells into a nude mouse. The procedure aims to infuse the cells into the left cardiac ventricle to disseminate the cells throughout the entire body (*left panel*). However, the needle may be inadvertently inserted in the right ventricle, resulting in migration and trapping of the cell in the lungs (*middle panel*). In other cases, the needle may miss the target entirely, resulting in cells being injected into the pericardial cavity (*right panel*). PET and CT images were visualized with OsiriX software as maximum intensity projections, with a slice thickness of 21.2 mm.

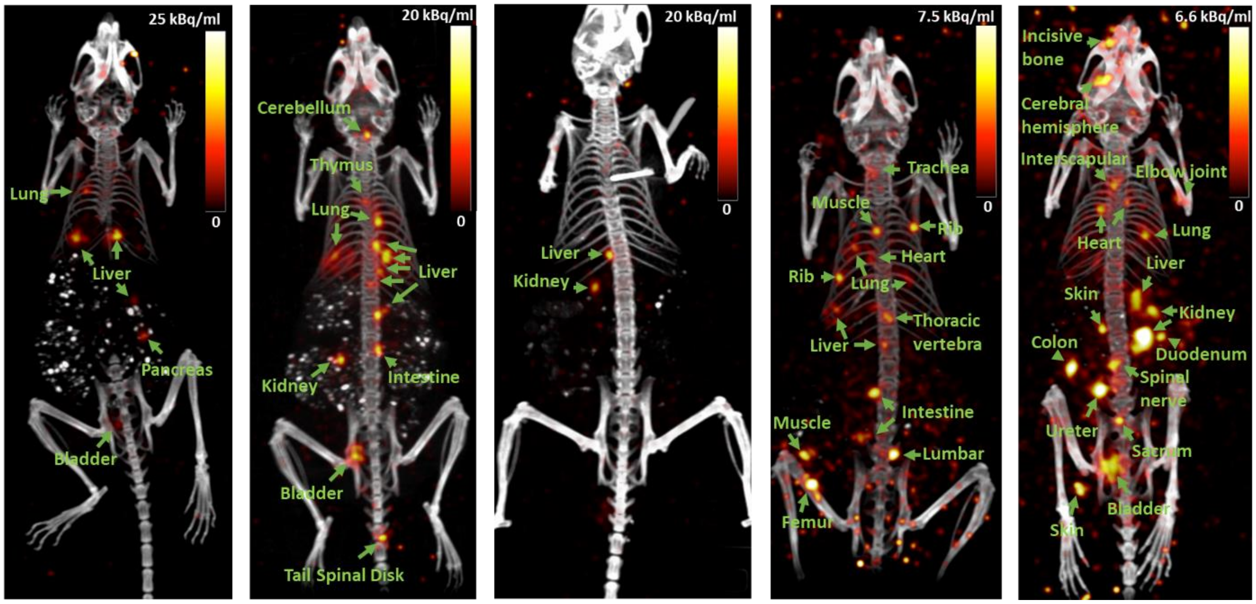


**Supplementary Fig. 9. Single-cell distribution in 5 mice following intracardiac injections.** PET/CT images showing the distribution of FDG-labeled B16 cells after intracardiac injection in five mice (in addition to the one shown in **Fig. 4**). The number of cells detected in each mouse varied from two to twenty, and the positions of these cells were categorized by a board-certified orthopedic surgeon (coauthor YW) by inspecting the PET/CT slices using the OsiriX software. PET and CT images shown above are displayed as maximum intensity projections using OsiriX software, with a slice thickness of 21.2 mm to display cells anywhere within the mouse.

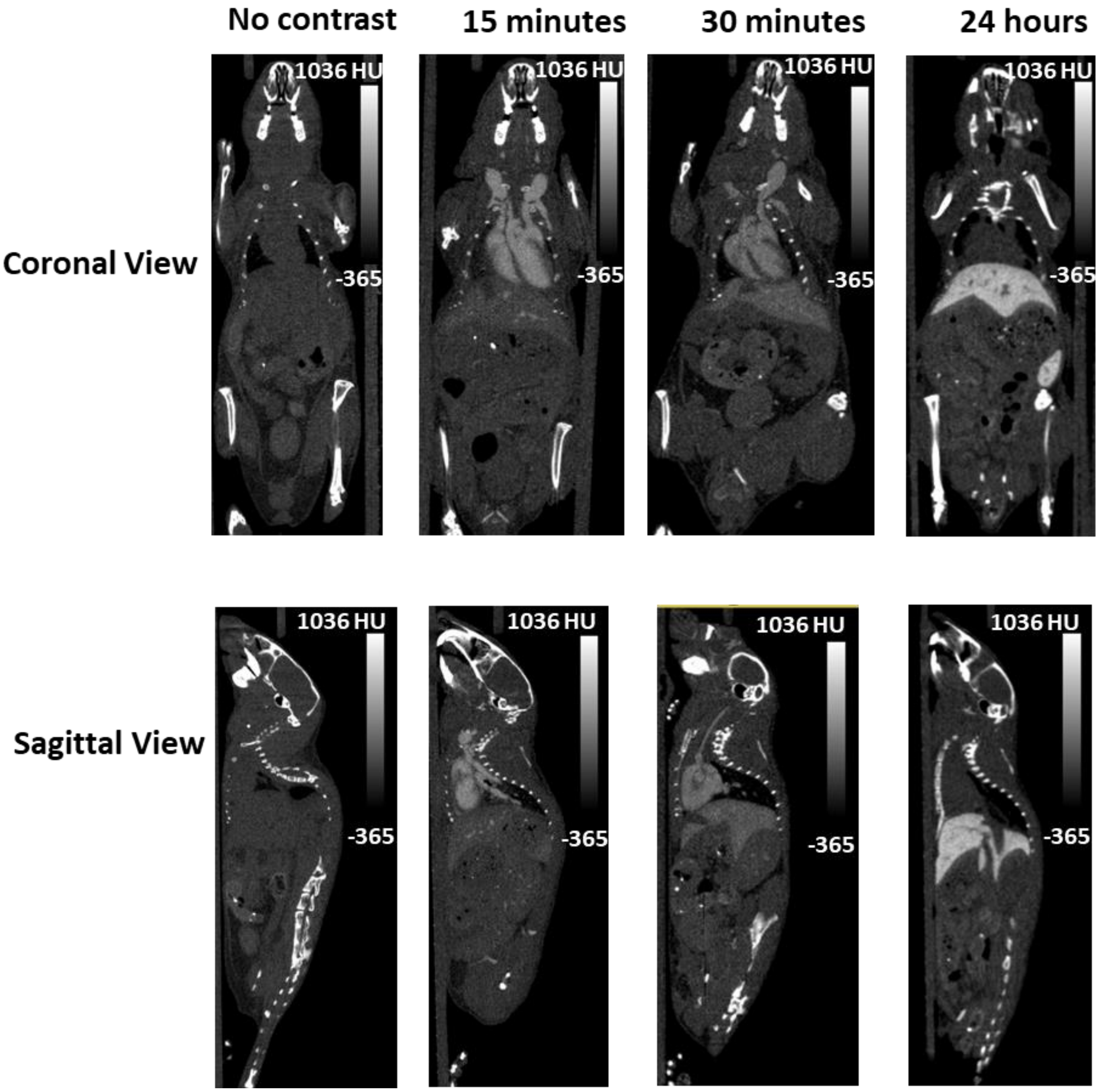


**Supplementary Fig. S10. Contrast-enhanced CT imaging of mouse vasculature.** CT imaging of a nude mouse after receiving an IV injection of a blood-pool enhancing contrast agent (ExiTron nano 12000). CT imaging was performed on a Sofie GNext scanner (80 kVp, bin 1, 100 μm resolution) at 15, 30, and 24 hours post-injection. Coronal and sagittal views of the mouse revealed that the concentration of the contrast agent peaked in the bloodstream within 15 minutes. The signal was dimmer at 30 minutes due to liver clearance. At 24 hours, the contrast enhancement was localized in the liver and spleen.

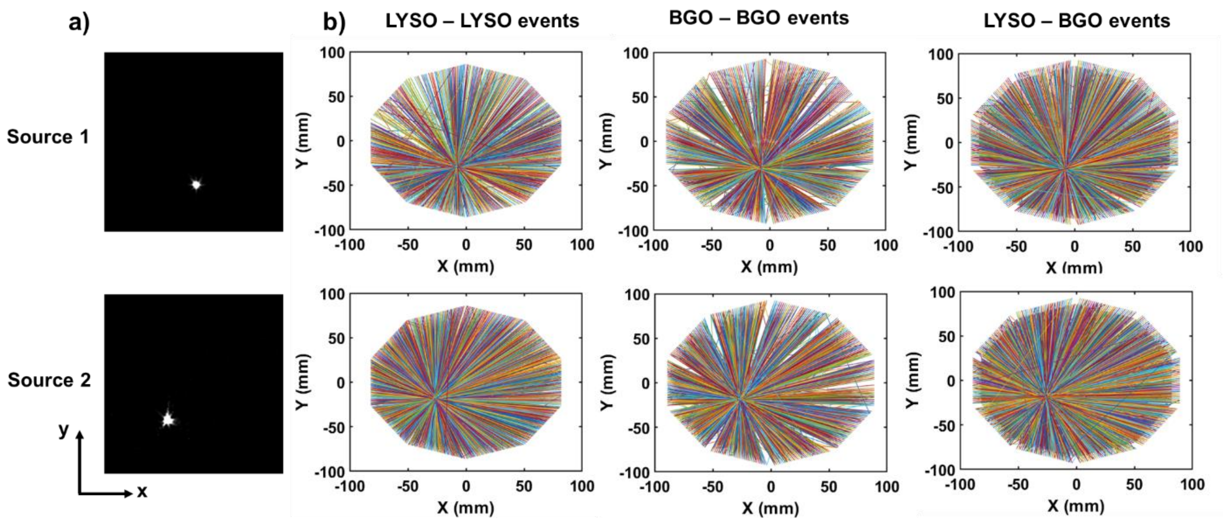


**Supplementary Fig. 11. Validation of list-mode import tool.** A custom software tool was developed to import the list-mode data packets generated by the scanner into a human-readable format suitable for reconstruction. **(a)** Standard OSEM-reconstructed images of two $^{22}Na$ point sources, located in two different axial planes ($Z_1$ = 41.3 mm and $Z_2$ = 92.4 mm), containing 66.6 kBq and 222 kBq of radioactivity, respectively, **(b)** Axial plane plots showing list-mode coincidence events (colored lines), which were recorded by the scanner and converted using the software tool. Coincidence events between the LYSO and BGO ring within the scanner are shown on separate plots. The lines corresponding to these events converged to a location consistent with the positions of the point sources generated using the vendor-provided OSEM image reconstruction software, indicating that the conversion tool correctly imported the list-mode data into MATLAB.

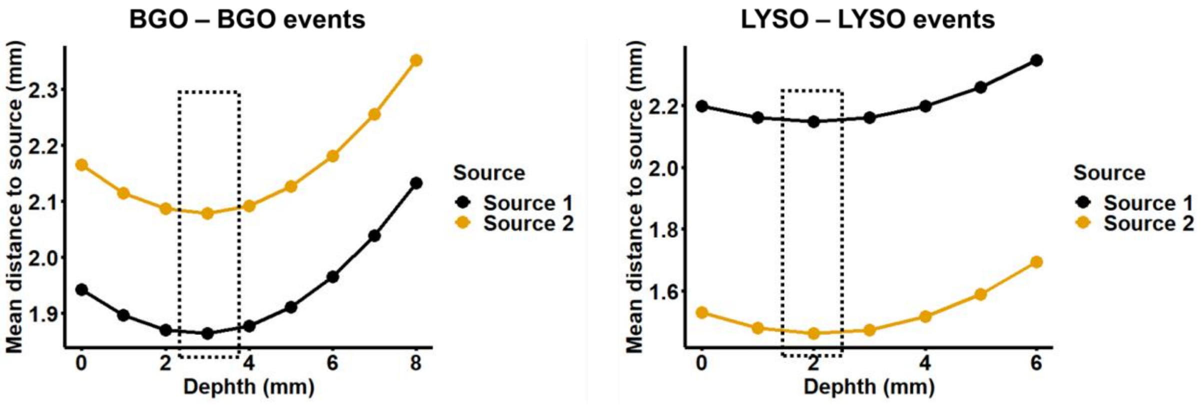


**Supplementary Fig. 12. Depth of interaction optimization within scintillator crystal.** PET detectors are not able to measure the depth of interaction of the annihilation photon within each scintillation crystal. Here, we chose the photon interaction depth such that the mean distance between the coincidence events recorded in the axial plane and the estimated source location is minimized. The analysis shown above shows that the optimal depth of interaction for BGO and LYSO crystals is around one-third of the crystal length, or 3 mm and 2 mm, respectively. This value was used to convert the machine data into accurate spatial coordinates.

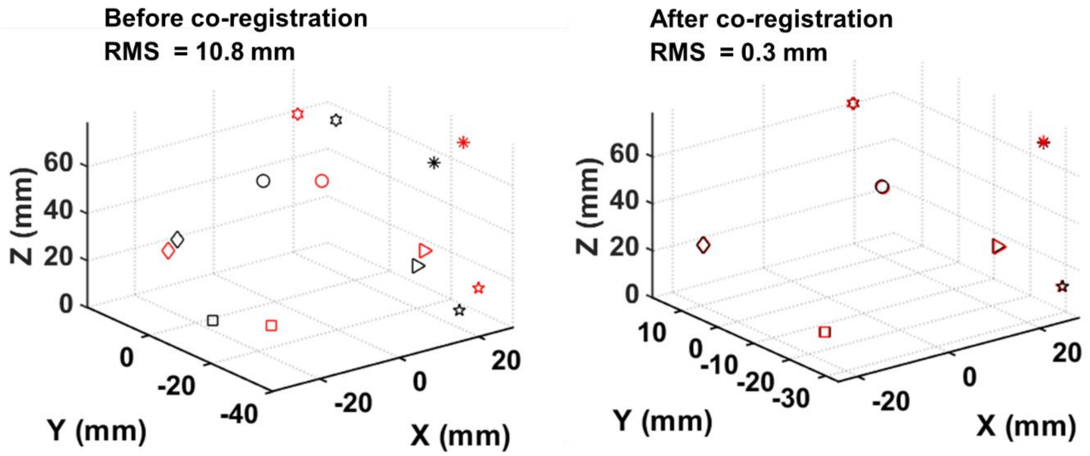


**Supplementary Fig. 13. Spatial co-registration of converted list-mode data with vendor-provided PET-CT coordinates.** The 3D positions of seven $^{22}Na$ point sources (activity between 66.6 kBq and 222 kBq) were estimated after reconstructing the list-mode data using our custom PEPT-EM method (black labels) and the vendor-provided OSEM image reconstruction (red labels). As shown in the left panel, the two methods yield slightly different locations due to the use of different reference coordinate systems. To resolve this issue, we used an affine transformation to co-register the locations of the seven-point sources estimated by these two methods (right panel). The transformation reduced the difference between the two methods from 10.8 mm to 0.3 mm.

**Supplementary Methods**

**A. List-mode import tool**

The first part of the supplementary method explains the steps required to i) extract the line-of-response (LOR) coordinates and time stamps from the list mode data (.lmf file) generated by the Sofie GNext software, ii) Optimize the annihilation photon penetration depth, and iii) Co-register the converted PET data to the output of the OSEM reconstruction software provided by the vendor.

1. **List mode to LOR data conversion**
   The Sofie Gnext PET scanner comprises 2 rings, each with 10 square detector panels.[1] Within each detector panel,  there are two layers of scintillators: BGO (36 x 36 crystals) and LYSO (48 x 48 crystals). We first extracted the crystal IDs and detector IDs from the .lmf file generated by Sofie GNext with the following structure. The crystal IDs and detector IDs were respectively packed in bits 41-52 & 53-64 (values 0-2303) and bits 23-27 & 28-32 (values 0 – 19) of the 64-bit lmf packet. We then made the following orientation assumptions and used geometry to convert the crystal IDs and detector IDs to spatial coordinates of detected annihilation photons:
   - The 10 detector panels within each detecting ring are oriented, as shown in the left figure below. The detector IDs are labeled clockwise, 1-10 in the first ring and 11-20 in the second ring.
   - The crystal IDs in each detector panel are indexed from left to right, top to bottom.
   - The annihilation photons are assigned to the center of each crystal.

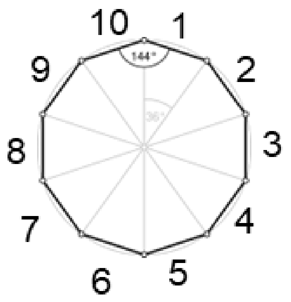


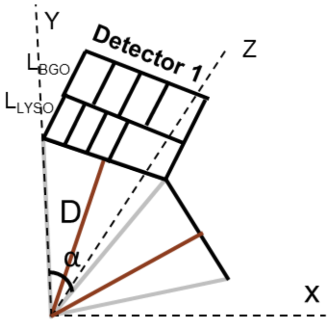


The 3D coordinates ($X, Y, Z$ ) representing the location of detected annihilation photons for the BGO or LYSO detector panels were calculated as follows.

$$X = x \times \text{pitchsize} \times \cos\left(\frac{(2i-1)}{2}\alpha\right) + D\sin\left(\frac{(2i-1)}{2}\alpha\right)$$

$$Y = -x \times \text{pitchsize} \times \sin\left(\frac{(2i-1)}{2}\alpha\right) + D\cos\left(\frac{(2i-1)}{2}\alpha\right)$$

$$Z = y \times \text{pitchsize} + \frac{a}{2}$$

$$x = mod(ID/48) - 23.5 \qquad y = \text{floor}(ID/48) - 23.5 \qquad D = \frac{a}{2\tan\left(\frac{\alpha}{2}\right)}$$

where $ID$ is the crystal ID on each panel (values 0-2303 for LYSO and 0-1295 for BGO); $i$ is the detector ID (value 1-10); $a$ is the detection panel width ($a$ = 52 mm); $D$ is the distance between the center of the ring to the front of each panel; $pitchsize$ is the crystal pitch (value of 1.09 mm for LYSO and 1.63 mm for BGO); $\alpha$ is the angle from the center of the ring between two adjacent panels, as shown in the right panel ($\alpha = 36^0$). A LOR is formed by connecting two coincident (simultaneously detected) annihilation photons events.

2. **Penetration depth optimization**

The LYSO and BGO crystals are 6.1 mm and 8.9 mm long, respectively. In the formulas used to calculate the 3D coordinates ($X$, $Y$, $Z$ ) derived in section 1, we assumed the annihilation photons were absorbed in the surface of the first layer of the scintillator (LYSO). In reality, these photons travel through a certain depth of scintillator material before being absorbed. To determine the penetration depth of the annihilation photons within each crystal layer ($d_{LYSO}, d_{BGO}$), we first sampled the penetration depth term (highlighted in red) in the formulas calculating the 3D coordinates of the detected annihilation photons, as shown in the equations below. We then varied $d_{LYSO}$ and $d_{BGO}$ within the depth of each crystal layer ($d_{LYSO} \epsilon$ [0, 6.1], $d_{BGO} \epsilon$ [0, 8.9]) to find the depth at which the LORs converged closest to the true locations of the $^{22}$Na point sources (determined by OSEM constructed image). We observed that the optimal penetration depth for LYSO and BGO crystals was around one-third of the crystal length, or 2 mm and 3 mm, respectively (**Supplementary Fig. 12**).

The set of formulas used for the optimization of penetration depth in LYSO crystal is as follows:

$$X = x * pitchsize * \cos\left(\frac{(2i-1)}{2}\alpha\right) + D*sin\left(\frac{(2i-1)}{2}\alpha\right) + d_{LYSO} * sin\left(\frac{(2i-1)}{2}\alpha\right)$$

$$Y = -x * pitchsize * \sin\left(\frac{(2i-1)}{2}\alpha\right) + D*cos\left(\frac{(2i-1)}{2}\alpha\right) + d_{LYSO} * cos\left(\frac{(2i-1)}{2}\alpha\right)$$

$$Z = y * pitchsize + \frac{a}{2}$$

The set of formulas used for the optimization of penetration depth in BGO crystal is as follows:

$$X = x * pitchsize * \cos\left(\frac{(2i-1)}{2}\alpha\right) + D*sin\left(\frac{(2i-1)}{2}\alpha\right) + (L_{LYSO} + d_{BGO}) * sin\left(\frac{(2i-1)}{2}\alpha\right)$$

$$Y = -x * pitchsize * \sin\left(\frac{(2i-1)}{2}\alpha\right) + D*cos\left(\frac{(2i-1)}{2}\alpha\right) + (L_{LYSO} + d_{BGO}) * cos\left(\frac{(2i-1)}{2}\alpha\right)$$

$$Z = y * pitchsize + \frac{a}{2}$$

where $L_{LYSO}$ is the full length of the LYSO crystal ($L_{LYSO}$ = 6.1 mm).

3. **Co-registration of LOR data to Sofie CT/PET**

To co-register the LOR coordinates generated by the aforementioned list-mode conversion method to the output of the vendor software, we imaged seven $^{22}$Na point sources placed in the PET scanner field of view. We then applied an affine transformation (as shown below) to calculate the transformation matrix $M$ from the locations of point sources estimated by our list-mode conversion ($x\ y\ z$) and the locations estimated by the vendor reconstruction software ($x'\ y'\ z'$). The transformation reduced the difference between the locations estimated by the two methods from 10.8 mm to 0.3 mm (**Supplementary Fig. 13**). The transformation matrix (shown below) rotated the coordinates 16.26° in the axial plane and translated the coordinates 1.15 mm along the x-axis.

$$p' = Mp$$

$$\begin{pmatrix} x' \\ y' \\ z' \\ 1 \end{pmatrix} = \begin{pmatrix} a & b & c & t_x \\ d & e & f & t_y \\ g & h & i & t_z \\ 0 & 0 & 0 & 1 \end{pmatrix} \begin{pmatrix} x \\ y \\ z \\ 1 \end{pmatrix}$$

**Rotate 16.26° in xy plane** **Translate 1.15 mm in x axis**

$$M = \begin{pmatrix} 0.96 & -0.31 & 0 & 1.15 \\ 0.3 & 0.96 & 0 & 0 \\ 0 & 0 & 1 & 0 \\ 0 & 0 & 0 & 1 \end{pmatrix}$$

### B. PEPT-EM algorithm

The PEPT-EM tracking algorithm was originally reported by Manger et al. to track the 3D positions of chemical particles labeled with large amounts of radioactivity.[2] The algorithm employs an iterative approach, switching between clustering list-mode data and estimating new positions for the radioactive sources, ultimately converging towards the maximum likelihood positions of these sources. The iterative process begins by initializing the position $x_t$, variance $\sigma_t^2$, and activity $r_t$ of each of the K sources, followed by calculation of weights $w_{l,k}$, which represent the relative contribution of each LOR to each source. These weights are then used to compute a new estimate of the sources' positions, defined as the centroid of the weighted LORs. The variance and activity of each source are also updated during this step. The main steps of the algorithm are summarized in the following diagram, where $D(x_k, l)$ is the distance between a source $x_k$ and a LOR $l$, $r_0$ describe the contribution of background counts such as scattering and random coincidences, $\alpha$ is a user-defined parameter interpreted as the inverse variances of the background cluster, $I_3$ is the identity matrix, and $y_l$ and $\boldsymbol{n_l}$ characterize the length and direction vector of LOR $l$.

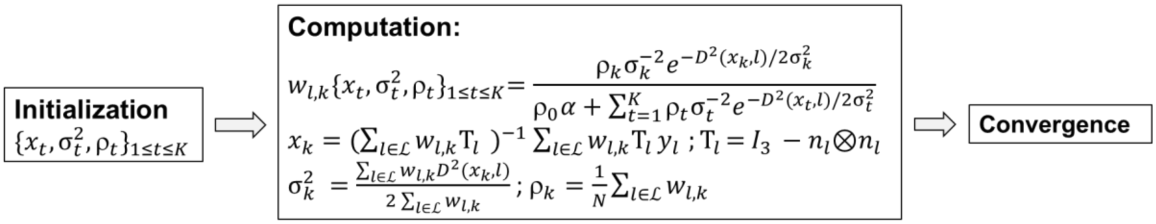

### C. Calculation of radiation dose received by cells during FDG labeling

The radiation dose received by cells while being labeled with FDG was estimated based on the total positron energy deposited in the media volume. We first calculated the number of decay (or the number of positrons generated) that occurred during the labeling as the difference between the number of radionuclides at time 0 ($N_0$) and time $t$ = 60 minutes ($N$):

$$N_0 - N = N_0 - N_0 e^{-\lambda t} = N_0(1- e^{-\lambda t}), \qquad \lambda = \frac{ln2}{t_{1/2}}\ , \qquad N_0 = \frac{A}{\lambda}$$

where $t_{1/2}$ is the half-life of $^{18}$F ($t_{1/2}$ = 109.8 minutes), and $A$ is the radioactivity at time 0 ($A$ = 740 MBq).

The radiation dose $S$ deposited in the cell culture media volume is estimated as the total positron energy (average energy of one positron is $e_p$ = 250 keV) over the mass of the media ($m$ = 2 g). Accordingly, the deposited dose was

$$S = \frac{(N_0 - N)e_p}{m} = \frac{N_0(1- e^{-\lambda t})e_p}{m} = \frac{\frac{20\ mCi \times 3.7 \times 10^7 \left(\frac{Bq}{mCi}\right)}{0.000105(s^{-1})}\left(1-e^{-(0.000105\ \times 3600)}\right) \times 250\ keV \times 1.6 \times 10^{-16}\ \left(\frac{J}{keV}\right)}{0.002\ kg}$$

$$= 44\ \text{Gy}$$


### Supplementary Video Captions

**Video S1.** Evolution of the cell position estimate during the convergence of the PEPT-EM algorithm, shown for 5 single cells in vials (4-32 Bq) and displayed over the matching CT image. The algorithm was initialized assuming random cell locations. As the algorithm iterates, it progressively clusters the list-mode data and converges towards the maximum-likelihood location of the cells, leading to a significant reduction in the standard deviation of the estimate (represented by the radius of each circle).

**Video S2.** 3D rendering of FDG-labeled B16 cells introduced into a mouse via intravenous injection. All the cells are arrested in the lung while traversing the network of small capillaries. The images were rendered using OsriX software.

**Video S3.** 3D rendering of FDG-labeled B16 cells introduced into a mouse via intracardiac injection (left ventricle). The cells were widely disseminated throughout the mouse body, arresting in various organs. The images were rendered using OsriX software.

**Video S4.** Evolution of the cell position estimate during the convergence of the PEPT-EM algorithm, shown for 14 single cells administered into a mouse via intracardiac injection and displayed over matching CT image (sagittal view). The algorithm was initialized by assuming random cell locations near the center of the field of view. As the algorithm iterates, it progressively clusters the list-mode data and converges towards the maximum-likelihood location of the cells, leading to a significant reduction in the standard deviation of the position estimate (represented by the radius of each circle).

**Video S5.** Mouse cardiac left ventricle imaged with ultrasound (Vevo 2100, Visual Sonics, 550S probe) in B-mode (40 Hz) as guidance for intracardiac injection. The transducer was oriented to image the short axis of the heart. The syringe needle appeared in the first three seconds during the needle insertion and injection (highlighted in the top left corner of the video).

**Video S6.** 3D rendering of FDG-labeled B16 cells, which were introduced into a mouse via intracardiac injection. The cardiovascular and bony anatomy were segmented from the matching CT scan and are displayed alongside the PET image of the single cells. A blood-enhancing contrast agent was utilized to highlight large blood vessels, including the aorta and carotid artery, among others. Notably, the cancer cells were not detected in the vicinity of these major vessels; instead, they appeared to arrest in the network of tissue capillaries, which cannot be imaged by CT due to the limited resolution of 100 μm.
